# China Autism Brain Imaging Consortium: Charting Brain Growth in Chinese Children with Autism

**DOI:** 10.1101/2025.02.20.639044

**Authors:** Lei Li, Miaoshui Bai, Kelong Cai, Doudou Cao, Xuan Cao, Jie Chen, Xue-Ru Fan, Peng Gao, Wenjing Gao, Dongzhi He, Fanchao Meng, Xi Jiang, Litong Ni, Xiuhong Li, Lizi Lin, Yingqiang Liu, Zhimei Liu, Ning Pan, Qi Qi, Bin Qin, Xiaolong Shan, Xiaojing Shou, Longlun Wang, Miaoyan Wang, Xin Wang, Dandan Xu, Yin Xu, Yang Xue, Ting Yang, Yun Zhang, Jinhua Cai, Huafu Chen, Aiguo Chen, Feiyong Jia, Haoxiang Jiang, Jin Jing, Tingyu Li, Shijun Li, Wei Wang, Jia Wang, Lijie Wu, Xuntao Yin, Rong Zhang, Xi-Nian Zuo, China Autism Brain Imaging Consortium, Xujun Duan

**Affiliations:** The Clinical Hospital of Chengdu Brain Science Institute, School of Life Science and Technology, University of Electronic Science and Technology of China, Chengdu 610054, PR China; Department of Developmental and Behavioral Pediatrics, Children’s Medical Center, The First Hospital of Jilin University, Jilin University, Changchun 130021, PR China; College of Physical Education, Yangzhou University, Yangzhou 225127, PR China; School of Sport and Brain Health, Nanjing Sport Institute, Nanjing 210014, PR China; Department of Radiology, First Medical Center, Chinese PLA General Hospital, Beijing 100853, PR China; Department of Children’s and Adolescent Health, Public Health College of Harbin Medical University, Harbin 150086, PR China; Children’s Nutrition Research Center, Ministry of Education Key Laboratory of Child Development and Disorders, National Clinical Research Center for Child Health and Disorders, China International Science and Technology Cooperation Base of Child Development and Critical Disorders, Children’s Hospital of Chongqing Medical University, Chongqing 400042, PR China; State Key Laboratory of Cognitive Neuroscience and Learning, Developmental Population Neuroscience Research Center at IDG/McGovern Institute for Brain Research, Beijing Normal University, Beijing 100875, PR China; Department of Radiology, Guangzhou Key Laboratory of Child Neurodevelopment, Women and Children’s Medical Center Affiliated to Guangzhou Medical University, Guangzhou 510623, PR China; Department of Neurobiology, School of Basic Medical Sciences, Peking University Health Science Center, Beijing 100191, PR China; The National Clinical Research Center for Mental Disorders & Beijing Key Laboratory of Mental Disorders, Beijing Anding Hospital, Capital Medical University, Beijing, PR China; School of Public Health, Shenzhen, Sun Yat-sen University, 66 Gongchang Road, Guangming District 518107, Shenzhen, PR China; Department of Maternal and Child Health, Joint International Research Laboratory of Environment and Health, Ministry of Education, Guangdong Provincial Engineering Technology Research Center of Environmental Pollution and Health Risk Assessment, School of Public Health, Sun Yat-sen University, Guangzhou 510080, PR China; Department of Rehabilitation Medicine, Affiliated Hospital of Yangzhou University, Yangzhou 225001, PR China; Key Laboratory of Brain, Cognition and Education Sciences, Ministry of Education, Institute for Brain Research and Rehabilitation, Guangdong Key Laboratory of Mental Health and Cognitive Science, South China Normal University, Guangzhou 510630, PR China; Department of Radiology, Children’s Hospital of Chongqing Medical University, Chongqing 400042, PR China; Key Laboratory for Neuroscience, Ministry of Education/National Health Commission, Peking University, Beijing 100191, PR China; Department of Radiology, Affiliated Children’s Hospital of Jiangnan University, Wuxi 214000, PR China; Nanjing Sport Institute, Nanjing 210014, PR China; Department of Radiology, Affiliated Hospital of Yangzhou University, Yangzhou 225001, PR China

**Author notes:** co-corresponding authors of this work.

## Abstract

Autism Spectrum Disorder (ASD) is a lifelong neurodevelopmental condition characterized by atypical brain growth. While advances in neuroimaging and openly sharing large-sample datasets such as the Autism Brain Imaging Data Exchange (ABIDE) have improved understanding of ASD, most studies focus on adolescents and adults, with early brain development-critical for diagnosis and intervention-remaining underexplored. Existing research predominantly involves Western samples, offering limited insight and generalizability into non-Caucasian populations. We introduce the China Autism Brain Imaging Consortium (CABIC) (https://php.bdnilab.com/resources/), a grassroots effort by researchers across the country to aggregate previously collected multi-site structural MRI datasets and phenotypic information from 1,451 autistic children and 1,119 typically developing children, covering an age range from early childhood to school age (1.0 - 12.92 years). Here, we present this resource and depict brain growth charts to push forward a more comprehensive understanding of the brain development in Chinese autism children. We constructed brain growth charts that reveal a developmental shift in autistic children, transitioning from early overgrowth to delayed maturation. Regional analyses identified distinct atypical trajectories across specific brain regions. Individual deviation scores quantified inter-subject variability, characterizing the heterogeneity of brain development in ASD. Comparative analyses between CABIC and ABIDE highlighted differences potentially attributable to ethnicity and culture, advancing our understanding of cross-population neurodevelopmental diversity. CABIC MRI datasets will be shared publicly to foster investigation of the potential neural mechanisms underlying ASD in non-Western populations and support efforts toward precision medicine for autistic individuals across diverse backgrounds.

## Introduction

Autism spectrum disorder (ASD) is a childhood-onset neurodevelopmental condition characterized by social-communicative difficulties, restrictive and repetitive behaviors, and altered sensory processing in response to external stimuli ^1,2^. Over the past few decades, the global prevalence of ASD has risen dramatically, attributed to a confluence of factors, including heightened public awareness, advancements in diagnostic methodologies, broadened diagnostic criteria, and the potential influence of genetic and environmental risk factors^3^. By 2020, data from the Autism and Developmental Disabilities Monitoring Network revealed a striking prevalence of 1 in 36 children aged 8 years in the United States, marking a paradigm shift in recognizing ASD as a significant public health concern^4^. This sharp increase in prevalence underscores the expanding burden on healthcare systems and highlights the urgent need for multidisciplinary strategies to address the challenges associated with ASD. Despite advances in public health initiatives and an expanding body of research, fundamental gaps persist in understanding the neural mechanisms underlying ASD. The concept of standardized growth charts for quantifying age-related changes was first introduced in the late eighteenth century and has since become a cornerstone of pediatric healthcare^5^. These charts provide invaluable benchmarks for assessing individual developmental trajectories against population norms, enabling early detection of atypical growth patterns^5,6^. Building on this foundational principle, the Lifespan Brain Chart Consortium (LBCC) recently established a comprehensive framework for mapping nonlinear trajectories of human brain morphology across the lifespan^7^. Leveraging large-scale structural MRI datasets, this initiative uncovered previously unreported neurodevelopmental milestones, offering profound insights into brain maturation and aging processes. Such a chart would provide a standardized framework for quantifying brain structural alterations in mental disorder, enabling precise tracking of its atypical neurodevelopmental trajectory. However, a critical gap remains in the development of a brain chart specific to ASD. Previous research has consistently demonstrated that ASD is associated with atypical cortical and subcortical developmental patterns ^8–10^. Notably, ASD appears to follow a distinctive trajectory of brain maturation characterized by early brain overgrowth in childhood, an accelerated decline during adolescence, and potential degeneration in young adulthood^11^. Longitudinal and cross-sectional neuroimaging studies further reveal age-related abnormalities, underscoring the developmental nature of these changes^9^. These findings emphasize the need to investigate the neural etiology of ASD from a developmental perspective. Understanding the atypical developmental trajectories of ASD in early life is essential for elucidating its underlying neural mechanisms and for advancing early diagnostic and therapeutic strategies.

A large sample of data covering different ages is an important prerequisite for exploring the underlying neural developmental mechanisms of ASD. Efforts by large-scale multi-site dataset consortium, such as the Autism Brain Imaging Data Exchange (ABIDE) and the Enhancing Neuroimaging Genetics Through Meta-Analysis (ENIGMA) ASD working group, have aggregated and shared extensive neuroimaging datasets from ASDs and matched typically developing children (TDC). These efforts have greatly advanced ASD brain imaging researches based on large-sample data and enhanced our understanding of ASD neurobiology. However, ASD is an early-onset neurodevelopmental condition, and a potential limitation of using the ABIDE and ENIGMA datasets for early-stage neurodevelopmental studies is the age range of the samples^12^. Early childhood is widely recognized as a crucial period for brain development due to the intense formation and fine-tuning of neural circuits ^13^. Depicting the neuroanatomical development of ASD in early childhood may help understand the progression in later stages of life and facilitate early diagnosis and intervention. However, previous studies have largely focused on measuring brain differences in autistic adolescents and adults. Early brain development at ages closer to clinical diagnosis and during critical periods for early intervention remains underexplored. Specifically, studies that integrate data across the full spectrum of early to late childhood and adolescence are still limited. Additionally, the majority of existing literature is based on Caucasian participants living in Western cultures. Consequently, it may not be possible to extrapolate the findings to individuals from different ethnic and cultural backgrounds, such as Chinese autistic children.

To address these issues, we introduce the China Autism Brain Imaging Consortium (CABIC) (CABIC: https://php.bdnilab.com/resources/; Chinese Color Nest Project: https://ccnp.scidb.cn/en), a grassroots effort by researchers across the country to aggregate previously collected multi-site structural MRI datasets and phenotypic information on 1,451 autistic children and 1,119 TDC, covering an age range from early childhood to school age. Our objective is to delineate the role of large-scale Chinese samples in investigating atypical developmental trajectories of ASD and in advancing the broader understanding of this complex neurodevelopmental condition. Another consortium’s objective is to identify and address heterogeneities that vary across studies, whether these differences are intentional or inadvertent. Recognizing and understanding these variances are crucial for guiding future research endeavors towards greater harmonization among research groups. We also anticipate developing unified clinical assessment and scanning strategies to guide future efforts aimed at increasing harmonization among research groups. This process is essential to promote consistency and comparability of findings across datasets, which will ultimately enhance the reliability and generalizability of research findings in the field of ASD neuroimaging.

Here, we present this resource and depict brain growth charts throughout childhood in Chinese children with ASD. The non-linear developmental trajectories of brain morphology in Chinese children with or without ASD were delineated by generalized additive models for location, scale, and shape (GAMLSS), which are recommended by the World Health Organization. This approach has been recently applied for lifespan development of human brain morphology by the Lifespan Brain Chart Consortium (LBCC) (https://github.com/brainchart/lifespan). We began by characterizing the changes and rates of changes in global and regional morphometric phenotypes, uncovering the developmental shift from over-to delayed maturation in Chinese autistic children. Subsequently, we identified and differentiated the various categories of abnormal developmental trajectories in distinct brain regions. Next, we presented inter-subject similarity characterizations to quantify the heterogeneity of Chinese autistic children using individualized deviation scores. Finally, to quantitatively estimate the diversity in brain growth attributable to ethnicity and culture, we conducted multidimensional comparisons to explore the differences between CABIC and ABIDE.

Overall, the CABIC initiative aims to fill existing knowledge gaps and offer valuable insights into the unique aspects of brain development in Chinese autistic children, contributing to the global understanding of ASD and supporting the development of tailored interventions and support strategies for this population.

## Results

### Sample composition

Ten sites from the CABIC contributed previously collected datasets for 2570 individuals (1451 autistic children and 1119 TDCs), covering an age range from early childhood to school age (1.0 – 12.92 years) (**Figure 1a**). Marked variation in the contributed number and age range across samples from different sites was evident (**Figure 1b**), along with a vast predominance of males (**Figure 1c**). For the detailed acquisition parameters and demographics of the datasets, see **Supplementary Table S1**.

**Figure 1.**
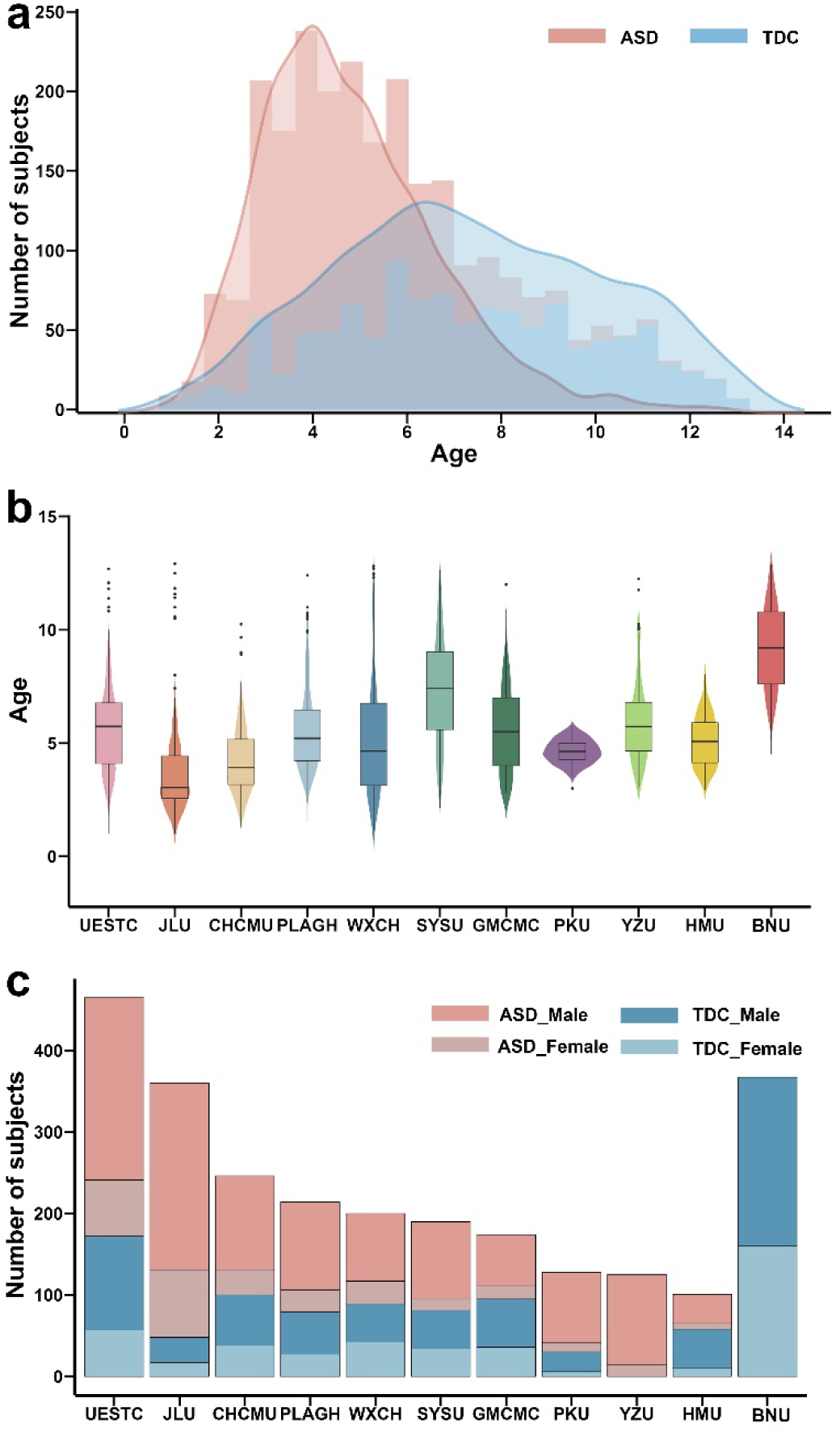
CABIC Sample Characteristics. **(a)** The distributions of age in ASD and TDC group **(b)** Age (in years) of participants per group for each contributing site. **(c)** Total number of males and females for each contributing site. Ordered by contributed sample number of per site.

All autistic children require a diagnosis of ASD by experienced clinicians, based on DSM-IV or DSM-5 criteria. Each site obtained supplemented clinical scores with one or more assessments, including the Childhood Autism Rating Scale (CARS, n = 395), the Autism Diagnostic Observation Schedule (ADOS, n = 476), the Autism Diagnostic Interview-Revised (ADI-R, n = 145), autism behavior checklist (ABC, 538), the Social Responsiveness Scale (SRS, n = 457), and/or Repetitive Behavior Scale-Revised (RBS-R, n = 514). Site-specific details for further information on diagnostics and assessments are available at https://php.bdnilab.com/sites/.

### Brain growth chart in Chinese autistic children

Normative growth charts for height, weight and head circumference are the cornerstone of pediatric health care. Analogously, normative models have recently been applied on brain structural and functional MRI phenotypes to elucidate the age-related nonlinear trajectories for healthy populations ^7,14–17^. In the current study, we constructed the brain growth chart of Chinese autistic children using GAMLSS, which offers a robust and flexible framework for modeling non-linear growth trajectories ^7,18^. We followed the approach and model estimation procedure of LBCC to estimate non-linear age-related trajectories stratified by sex from early childhood to school age. Specifically, GAMLSS models were fitted to structural MRI data for the five morphometric phenotypes (total gray matter volume [GMV], subcortical gray matter volume [sGMV], cerebral white matter volume [WMV], mean cortical thickness [MT], and surface area [SA]) in the samples of Chinese autistic children and Chinese TDC, respectively. Then, the first derivatives of the trajectories were calculated to evaluate the rate of change/velocity and inflection points. Notably, given the vast predominance of males in ASD, only the results of male participants are shown in main text, as brain charts are modelled sex-specifically. The detailed image quality control, model specification and estimation are presented in the **Methods**.

To provide basic developmental insight into the overall tissue volumes of the cerebrum in Chinese autistic children, we first characterized the typical and atypical trajectories of global morphometric phenotype. The growth curves for males were in presented the **Figure 2a**. Consistent with the LBCC findings, the normative brain chart of GMV showed a nonlinear increase, peaking at 7.9 years, followed by a near-linear decrease. The normative curves of TDC indicated rapid increases in sGMV, WMV and SA, and declines in MT from early childhood to school age. These results were reconciling previous observations, particularly that cortical thickness increases during the perinatal period and declines during later development ^19,20^.

**Figure 2.**
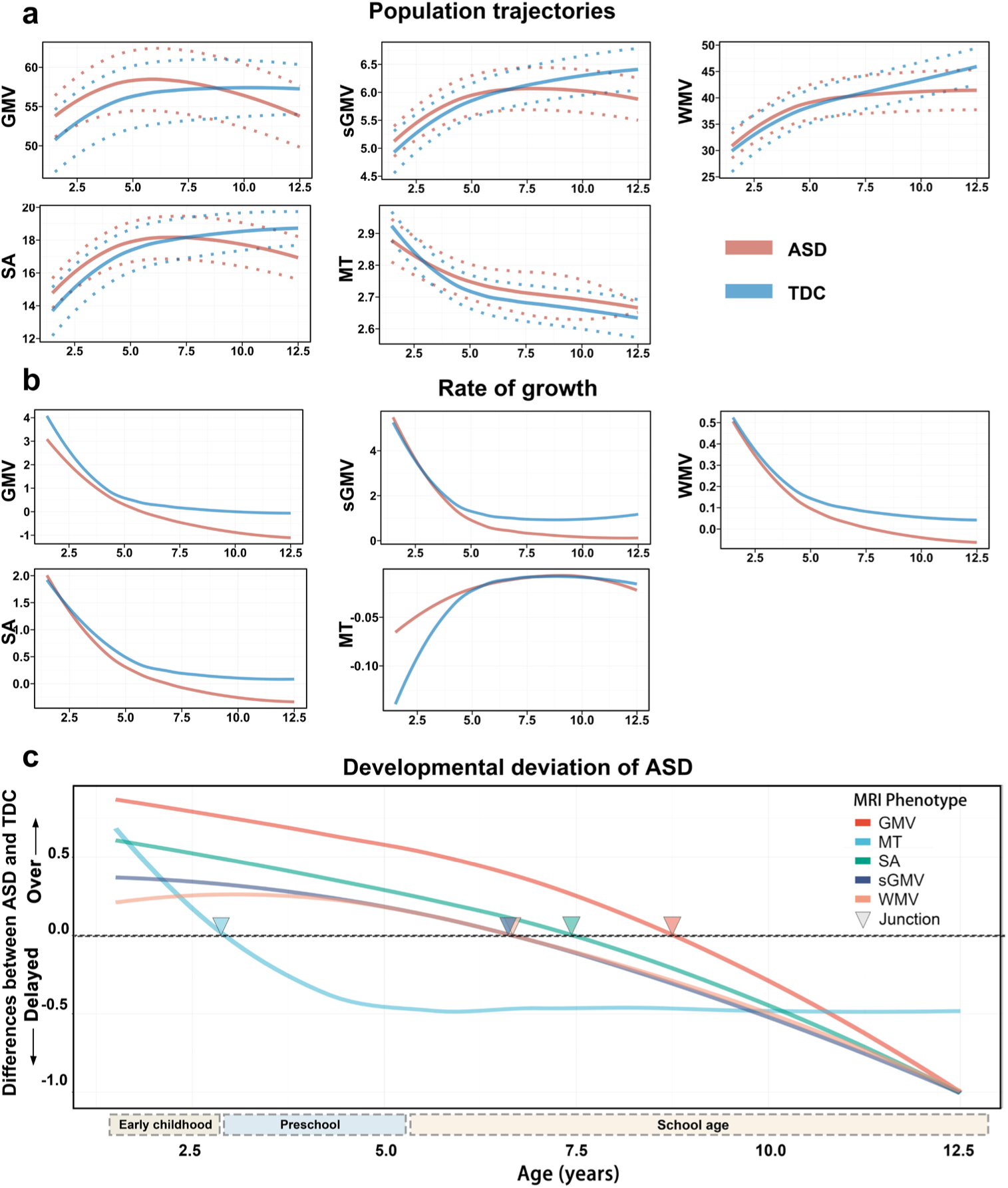
Typical and atypical whole-brain growth chart. **(a)** Left, typical and atypical trajectories of global morphometric phenotypes. The median (50% centile) is represented by a solid line, while the 2.5% and 97.5% centiles are indicated by dotted lines. Right, rate of growth (the first derivatives of the median trajectory). y axes are scaled in units of the corresponding MRI metrics (10,000 mm^3^ for volume value, 10,000 mm^2^ for surface area and mm for cortical thickness). **(b)** A graphical summary of the differences between the median (50th centile) in ASD and TDC group for each global MRI phenotype, and key developmental shift milestones. Triangles depict the key point of transition from overgrowth to delay in autistic children of each phenotype (defined by the zero of the differences of trajectory curves).

In terms of GMV, sGMV, WMV and SA, ASD curves were generally flatter and tilted downward compared to the TDC curves. During early childhood, the MRI metrics value was higher in the ASD group than in the TDC group. Then the ASD curve intersected with the TDC curve in school age (6-11y). However, instead of following the same pattern as TDC, the value for those with ASD kept decreasing through late childhood into adolescence. Notably, our results suggested that MT was smaller in ASD than TDC at age of 2 years. Subsequently, the ASD curve for MT was flatter compared to the TDC curve, with decelerated cortical thinning predominantly occurring throughout childhood. In addition, abnormal trajectories of growth rate in ASD appear to be conserved across morphometric phenotypes, characterized by accelerated growth earlier in early childhood, and accelerated decline throughout much of later childhood. In summary, the growth curves of global morphometric phenotypes showed an initial increase in early childhood, followed by an accelerated decline to meet the TDC curves, and subsequently continued to decline atypically into adolescence.

### Developmental shift milestone

As mentioned in the previous results, atypical brain growth charts in Chinese autistic children exhibited early childhood overgrowth, followed by accelerated decline and potential degeneration in late childhood. To further explore this pattern, we calculated the developmental difference curves (ASD-TDC) of the growth curves for each global morphometric phenotype. Developmental shift milestones were identified at the zero points of these difference curves (**Figure 2b**). These milestones mark the developmental shifted points from over to delayed maturation in autistic children compared to typically developing children. Among all global morphometric phenotypes, only MT showed a shift in early childhood (2.65 years), while GMV (8.76 years), sGMV (6.67 years), WMV (6.65 years), and SA (7.47 years) shifted during school age. Multi-dimension sensitivity analyses were performed for model stability and robustness **(Supplementary Information, Figure S1-S4)**.

### Extended regional cortical volumetric growth

To illustrate brain growth across regions, we averaged the regional GMV for participants in each age group (at 2-year intervals) and calculated the relative GMV compared to the maximum regional values (see Methods for details). The relative GMV for each age group is shown in **Figure 3a**. Our analysis revealed that different brain regions exhibited varying levels of relative maturity at different ages, suggesting that each region follows a distinct developmental trajectory. Thus, we extended the same GAMLSS modeling approach to estimate the regional volumetric growth chart for both ASD and TDC groups (**Figure S5**). Our findings revealed that the developmental trajectories of the majority of brain regions in Chinese autistic children parallel the atypical patterns observed in global morphometric phenotypes. Specifically, these trajectories are characterized by overgrowth in early childhood, followed by accelerated decline and potential degeneration in late childhood.

**Figure 3.**
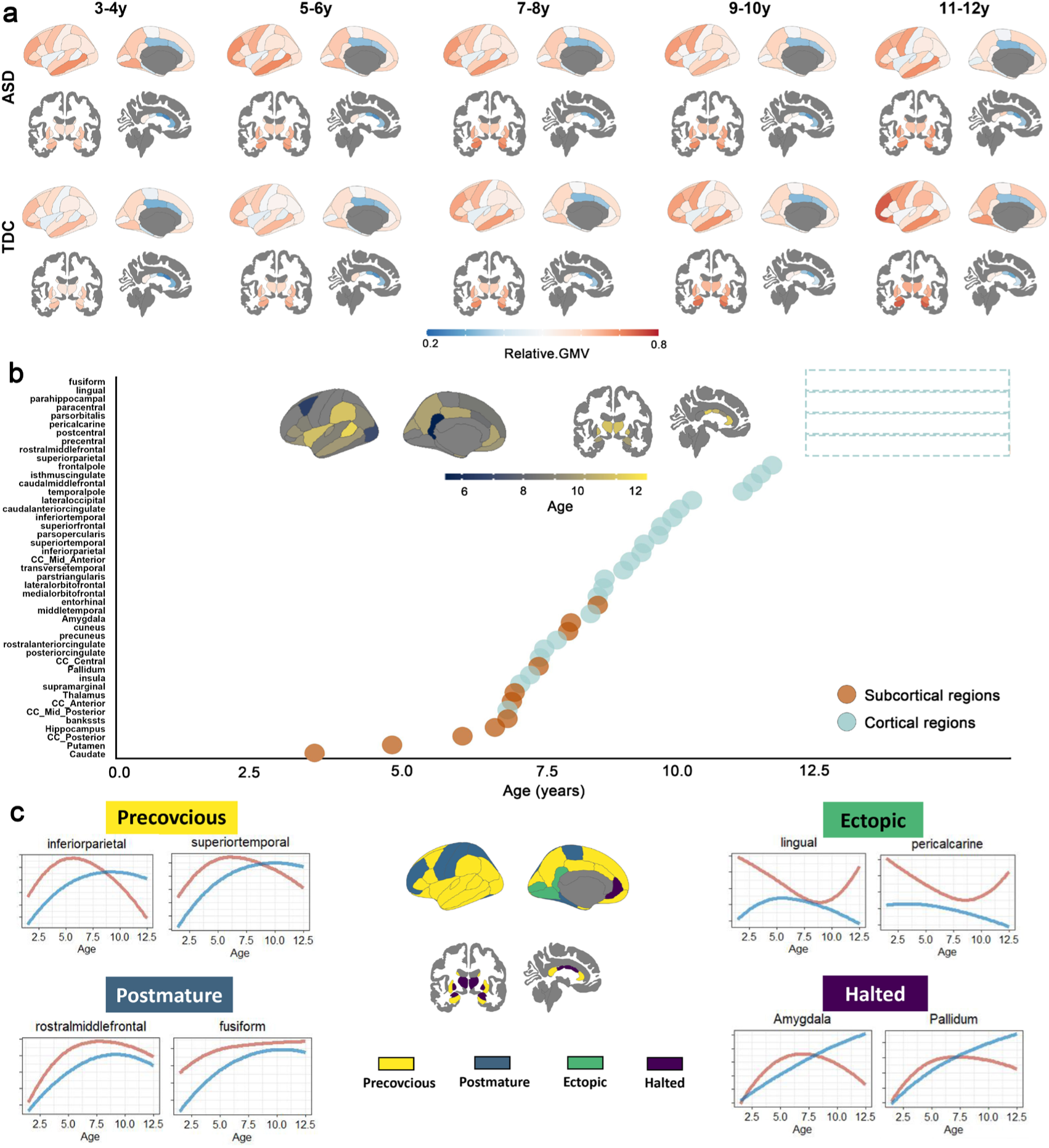
Regional brain growth chart of gray matter volume. **(a)** Regional relative value of gray matter volume compared to the regional maximum value for 34 bilateral brain regions, as defined by the Desikan–Killiany parcellation, and 12 subcortical regions. **(b)** A graphical summary of the regional developmental shift milestones for gray matter volumes. The dashed box indicates that these brain regions do not show a distinct milestone, i.e., there is no intersection in the developmental curves for autistic children and TDC. **(c)** Different categories of atypical regional developmental trajectories. Since models were generated from bilateral averages of each cortical region, the cortical maps are plotted on the left hemisphere purely for visualization purposes.

To analyze trajectories and milestones of brain development with finer-grained anatomical resolution, a similar series of methodological steps was performed for the regional GMV. Consistent with the results of global morphometric phenotypes, we found that the developmental shift milestones (from over to delayed maturation) of subcortical GMV generally occurred earlier than those of surface GMV (**Figure 3b**). Notably, some cortical regions did not show developmental shift milestones, suggesting that ASD and TDC growth curves in these regions did not exhibit intersection during this period.

### Incongruous regional developmental trajectories

As mentioned earlier, we found that the regional brain growth charts in autistic children exhibited inconsistent trends of atypical trajectories. As previously proposed by Martino et al., besides abnormalities in timing, changes in the shape of a trajectory may signal more profound developmental disturbances. Therefore, we employed a shape-based whole time-series clustering algorithm to identify different categories of atypical developmental trajectories across all brain regions in Chinese autistic children ^21^ (**Figure 3c**). We found that the association cortices, including most of the frontal and temporal cortices as well as the posterior parietal cortex (marked in yellow in **Figure 3c**), reflected precocious abnormality in developmental trajectories in ASD, relative to TDC. Specifically, GMV of those regions in ASD is larger than that in TDC during early childhood, and as development progresses, the GMV in TDC increases more rapidly and eventually surpasses that in ASD. The middle frontal cortex, fusiform and primary moto-soma cortex (marked in dark blue) reflected postmature abnormalities in developmental trajectories in ASD, relative to TDC, with the GMV in ASD being higher than that in TDC throughout childhood. The Inferior occipital cortex, involved lingual and calcarine sulcus (marked in green) reflected ectopic abnormality, characterized by atypical developmental changes in children with ASD that are not observed in TDC. In addition, extensive subcortical and limbic areas including amygdala, pallidum and middle cingulate gyrus, reflected halted abnormality, that is, the development of ASD ceases after initially following typical trajectories.

### Linking individualized deviation scores to behavioral variation

Recent literature has emphasized the potential of typical normative models to probe individual heterogeneity by enabling statistical inferences at the individual level ^7,22,23^. By benchmarking an individual’s brain measures against standards based on typically developing populations, this approach can quantify deviations from normative brain growth charts, providing unique insights into the typicality or atypicality of each individual’s brain structure. Therefore, in addition to depicting the atypical brain developmental trajectories of Chinese autistic children, we characterized the individualized deviation scores (age- and sex-specific) of each of their regional phenotypes with respect to the typical growth chart of the TDC (**Figure 4a**).

**Figure 4.**
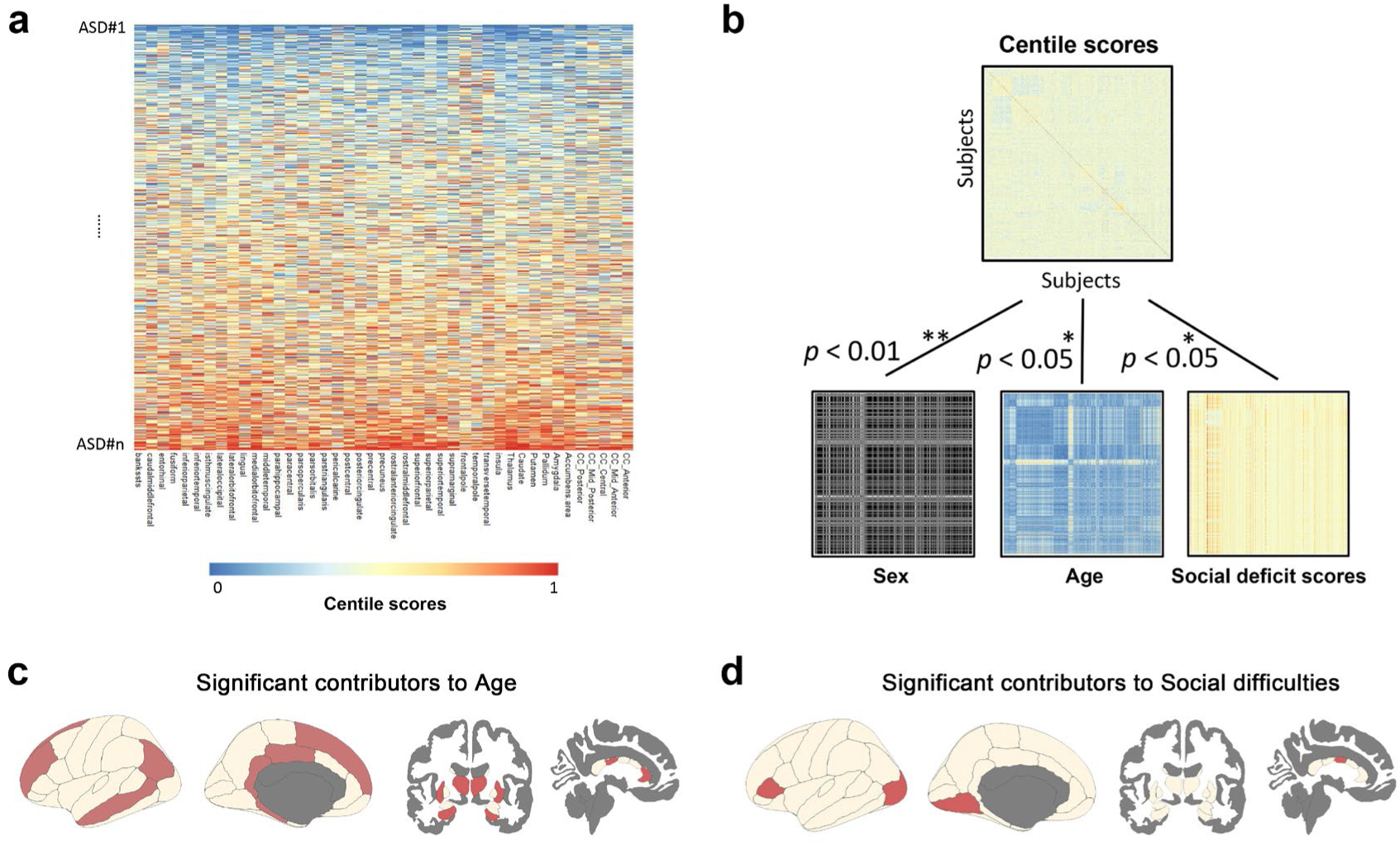
The relationship between individualized deviation scores of regional grey matter volume and phenotypic measurements. **(a)** Individual deviation centile scores for regional gray matter volume in all brain regions per autistic child. **(b)** The relationship between inter-subject similarity matrices of individualized regional grey matter volume deviation and different phenotypic measurements. **(c)** Brain areas that contributed significantly to age, based on Glmnet regression analysis. **(d)** Brain areas that contributed significantly to social cognition deficit scores, based on Glmnet regression analysis.

To test whether the individualized deviation scores correlate with individual variations in the behavioral and clinical characteristics of autistic children, representational similarity analysis (RSA) was used to obtain an inter-subjects similarity matrix which represents the similarity across participants with respect to the all-regional individualized deviation scores. This process was repeated and the similarity matrixes for each phenotypic measurement, including age, sex, site, scanning-site and social deficits score (SRS total scores) was calculated. By using Kendall rank correlation coefficient, our results indicated that there was significant correlation between similarity matrix of individualized deviation score and age (*τ* = 0.03, *p* < 0.001), sex (*τ* = 0.03, *p* < 0.001) and social deficits score (*τ* = 0.02, *p* < 0.001). There was no relationship with scanning-site (*τ* = 0.001, *p* = 0.780) (**Figure 4b**).

Then we used GLMnet-based multivariate sparse regression analyses to identify brain regions that drive the relationship between individualized deviation scores and age, sex, and social deficit scores^24^. Lambda traversal was employed to identify significant brain regions (**Figure S6**). Under the optimal model, only a few regions emerged as significant predictors, while all other brain regions had zero coefficients, indicating their lack of significance. We revealed 14 brain regions as significant predictors of age, including large portions of the cingulate gyrus and some temporoparietal regions (**Figure 4c**). Additionally, 4 brain regions were identified as significant predictors of social deficit scores, including the middle cingulate gyrus, paratriangular cortex, lingual gyrus, and lateral occipital lobe (**Figure 4d**). No brain region was found significantly contribute to sex.

### Sex differences in brain growth chart

Sex differences are increasingly recognized for their critical role in brain development ^25^. In our GAMLSS modeling, sex-stratified approach was used for establishing brain growth trajectories. Supplementary **Figures S7-S8** present the global and regional growth chart for females. These curves provide profound insights into the similar and distinct sex-dependent effects on morphometric phenotypes. Generally, both males and females exhibited similar nonlinear trajectories across multiple MRI phenotypes, indicating parallel developmental trajectories (**Figure S9-S10**). However, notable differences were also observed. Specifically, consistent with prior literature, we found that morphometric values in males were higher than that in females across various imaging phenotypes ^25^. These findings highlight the necessity of considering sex as a critical variable in constructing neurodevelopmental curves.

### Ethnical differences in ASD: Out-of-sample replication using the ABIDE autism cohort

To illustrate the differences in the autistic growth curves between populations, we utilized ABIDE samples from individuals aged 5-18 years **(Figure S11)** to construct the growth curves using the GAMLSS model. Our analysis revealed that the growth curve shapes for TDC in the CABIC and ABIDE datasets were quite semblable, with the notable exception that GMV was larger in ABIDE dataset than in CABIC dataset (**Figure 5a**). This observation aligns with previous studies indicating that Western children generally have larger GMVs than Chinese children ^15^. However, the brain growth charts for ASDs showed greater variation between the CABIC and ABIDE datasets (**Figure 5b**, group-differences in average amplitude of growth curve). To quantitatively assess the extent of diversity in brain growth attributable to ethnicity, we computed the normalized variance (NV) across various MRI phenotypes. The NV serves as an indicator of similarity or difference in growth curve shapes: a small NV suggests that the two brain growth curves are quite similar in shape, whereas a large NV indicates substantial differences. It is clear that the smaller NV values for TDC demonstrated a nearly identical growth trajectory across the two ethnic samples. In contrast, the growth patterns of ASD with larger NV values exhibited greater diversity in shape across the two populations, potentially influenced by ethnic, language and cultural factors (**Figure 5c**).

**Figure 5.**
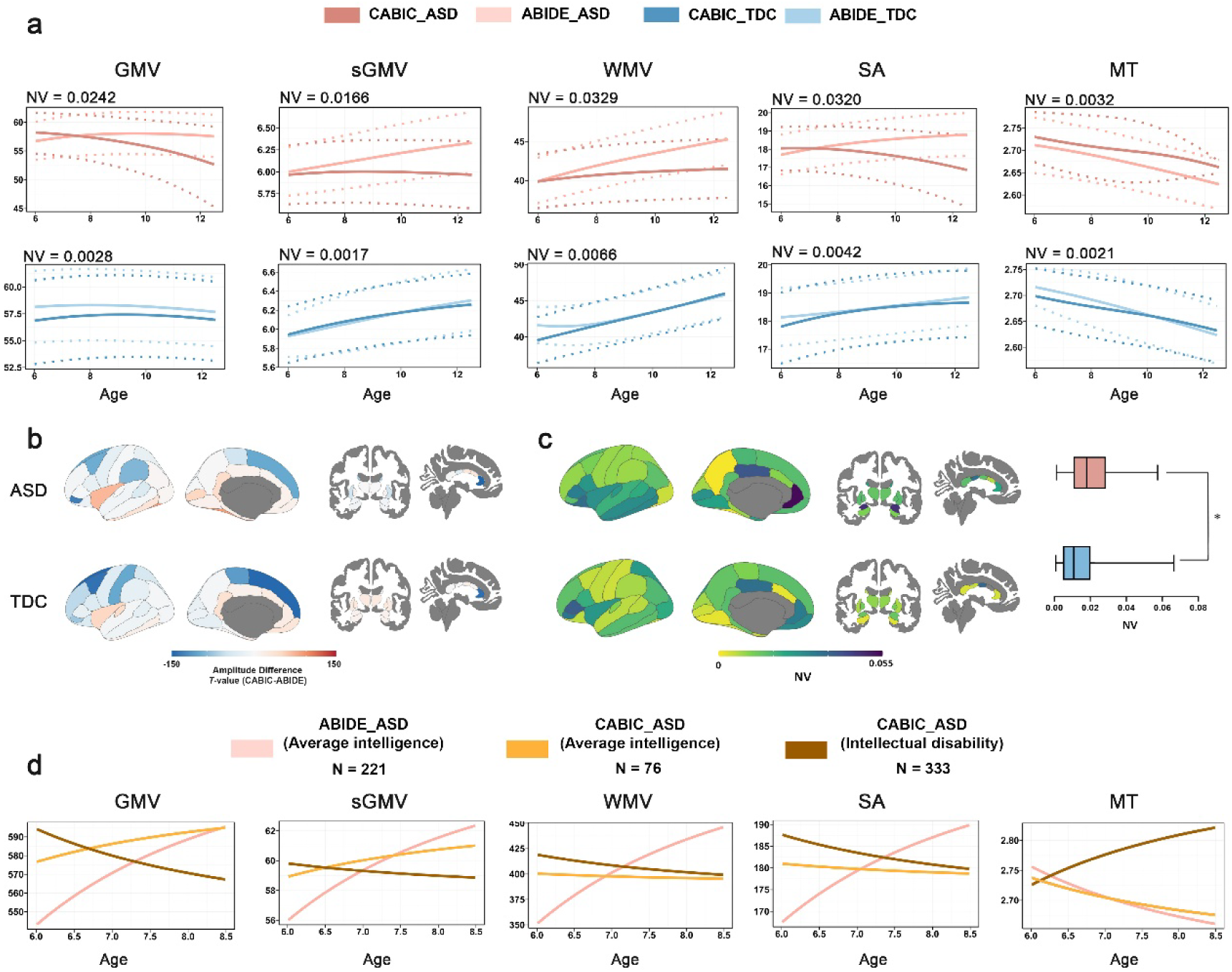
Similarities and differences in brain growth chart between CABIC and ABIDE. **(a)** Typical and atypical trajectories of global morphometric phenotype in CABIC and ABIDE. The median (50% centile) is represented by a solid line, while the 2.5% and 97.5% centiles are indicated by dotted lines. y axes are scaled in units of the corresponding MRI metrics (10,000 mm^3^ for volume value, 10,000 mm^2^ for surface area and mm for cortical thickness). **(b)** Group differences in amplitude of regional growth curves between CABIC and ABIDE **(c)** The regional NV values of the similarity between the CABIC and ABIDE in the ASD and TDC groups, respectively. Group difference in regional NV similarities across the whole brain between ASD and TDC was shown on the right. **(d)** Intelligence-stratified atypical brain growth chart of global morphometric phenotype in autistic children. (10,000 mm^3^ for volume value, 10,000 mm^2^ for surface area and mm for cortical thickness). See supplementary information for details in intelligence grouping. The left hemisphere is plotted here purely for visualization purposes. NV: normalized variance (NV).

Another potential factor influencing the developmental and clinical performance of ASD is intelligence ^26^. Full Scale Intelligence Quotient (FIQ) standard scores provided by ABIDE indicated significantly higher scores for TDC compared to ASD ^12^. In the inclusion samples of the current study, despite site-specific variations in the minimum FIQ, 97% of the datasets reported FIQ scores above 80. However, within the CABIC sample, only 18% of autistic children who underwent intelligence assessments had above-average intelligence (N = 76), while the remaining children exhibited intellectual disabilities (N = 333) **(See more details in Supplementary Information)**. To further investigate intelligence-specific growth patterns, we subdivided the CABIC autistic children into two groups based on their intelligence levels: those with average intelligence and those with intellectual disabilities. Using multivariable fractional polynomial regression (MFPR), we generated population- and intelligence-specific models. Our analysis revealed that within the CABIC dataset, both groups—autistic children with average intelligence and those with intellectual disabilities—exhibited similar growth trajectories in WMV and SA. In contrast, the ABIDE dataset showed almost opposite developmental patterns for these MRI phenotypes (**Figure 5d**). These findings suggest that the primary factors influencing WMV and SA growth in autistic children may be related to ethnicity or cultural differences. Additionally, our results showed that autistic children who have average intelligence in both the CABIC and ABIDE datasets exhibited similar growth trajectories in terms of GMV, sGMV and MT. Conversely, ASD with intellectual disabilities in the CABIC dataset demonstrated nearly opposite developmental patterns for these MRI phenotypes. These results indicated that intelligence may be a significant factor influencing GMV, sGMV, and MT growth in autistic children. Overall, these findings underscore the importance of considering both ethnic and intelligence-related factors when examining neurodevelopmental trajectories in ASD, highlighting the complex interplay of genetic, environmental, and cognitive influences in the neurodevelopmental trajectories of autistic children.

## Discussion

The establishment of the CABIC, which unites disparate and independent Chinese autistic research groups and aggregates previously collected MRI data, highlights the potential to accelerate advances in the field of neuroimaging in ASD through extensive collaboration. Our exploratory analysis of the CABIC dataset offers a comprehensive overview of the developmental brain growth trajectory in Chinese autistic children. Our results replicate and extend previously pivotal findings, and provides critical emerging themes that merit further study.

Standardized growth charts have long been integral to pediatric healthcare, providing benchmarks to assess developmental trajectories and identify atypical growth patterns. While the LBCC has developed a framework to map brain morphology across the human lifespan, no comparable chart exists for ASD. In the current study, we filled this gap by delineating the sex-stratified and age-related growth trajectories in multiple MRI-derived phenotypes. Developmental trends in current TDC group were generally consistent with those found in previous large-sample studies of typical brain growth chart ^7,27^. In addition, our findings align with previous autistic studies suggesting that ASD is a developmental disorder with atypical differences in whole and regional brain characteristics that evolve over time from early childhood into preadolescence. Our analysis of volumetric growth curves revealed a pattern of early brain overgrowth during the toddler years and early childhood in autistic boys and girls, followed by slowed growth in later childhood until the typical brain volume caught up with that of the autistic brain. These findings are consistent with previous studies that have reported increased head circumference and morphological volumetric overgrowth in autistic children during the early postnatal period ^28,29^. Furthermore, our results revealed that Chinese autistic children exhibit a thinner cortex at approximately 2 years of age, while experiencing slower cortical thinning throughout the following childhood. Overall, the atypical brain growth chart demonstrated that early postnatal life in autistic children is characterized by a relatively brief period, lasting several years or less, marked by abnormally accelerated brain overgrowth. This short phase is followed by a period of abnormally slowed, arrested, or degenerated growth between early childhood and preadolescence.

Our results supported the hypothesis of early brain overgrowth, one of the more widely reported neural phenotypes in ASD ^13,29,30^. Since Courchesne et al. first provided the evidence of atypical brain growth trajectory in ASD and propose the early brain overgrowth hypothesis ^8^, numerous neuroimaging studies have contributed to and refined the theory ^13,30^. This hypothesis accounts for inconsistencies in volumetric studies of ASD by postulating similar brain size at birth, followed by periods of accelerated growth over the next 2 years, and then deceleration of brain growth, equalizing volumes between groups by middle to late childhood. Mechanisms potentially underlying the early overgrowth in ASD may involve genetic, molecular, and cellular pathologies emerging during prenatal or very early postnatal life ^13^. The implications of early brain overgrowth are profound, as this period of rapid brain development coincides with critical windows for neurogenesis, synaptic pruning, myelination, and the establishment of neural networks ^31–33^. Rapid brain overgrowth can lead to an excessive number of neurons and synapses, resulting in atypical connectivity patterns hypothesized to contribute to the core symptoms of ASD. We speculate that the abnormally accelerated early brain overgrowth may implies innate abnormalities in initial neural and laminar morphological organization, as well as connectivity ^9^. These abnormalities are distinct from those influenced by experience and learning activities. In contrast, typical brain growth may reflect continuous refinement of brain organization through adaptive functional activity guided by experience and learning. Research in TDC suggests that slower growth in association cortex are associated with higher cognitive performance and better long-term outcomes ^34^. Notably, overgrowth and an excessive number of neurons will eventually trigger a late corrective or remodeling phase, involving attempts to prune the excessive number of abnormal axonal connections, synapses, and neurons to improve neural circuit function. The neuroinflammatory processes observed in ASD may reflect this secondary remodeling process, suggesting that the brain attempts to adapt and reorganize after an initial period of overgrowth ^35^. This corrective phase, while potentially beneficial, may also contribute to the atypical development observed in ASD.

Recent research suggests that although the anatomic pathology of autism is dynamic throughout development stage, early overgrowth is not ubiquitous across the brain, that is, some regions and structures display overgrowth while others do not ^36,37^. Our findings are consistent with previous findings suggesting that brain morphology changes throughout childhood, but the rates and types of change vary by brain region. Moreover, different ages likely exhibit specific atypicality. We observed that overgrowth in subcortical areas generally precedes that in cortical areas. Additionally, it is possible that some early-developing brain regions may experience abnormally accelerated overgrowth prior to the ages examined in our study. Thus, some regions did not show a developmental transition from over to delayed maturation. These highlight the importance of heterochronicity in understanding the phases of brain development in ASD.

Martino et al. noted that abnormalities in developmental timing and changes in trajectory shapes may indicate profound developmental disturbances ^38^. Our data further support this concept, identifying four categories of incongruous developmental trajectories in ASD. Specifically, most regional trajectories exhibited precocious or postmature abnormalities, consistent with early overgrowth observed in global MRI phenotypes. Additionally, extensive subcortical areas, including amygdala, pallidum and middle cingulate gyrus, demonstrated early overgrowth followed by halted development in later stages. Our recent clinical research demonstrated that personalized continuous theta-burst stimulation targeting the amygdala and its connected networks improved social and communication skills in minimally verbal children with ASD, potentially preventing the termination of neural growth and development ^39^. Accumulating neuropsychological evidence highlights those structural abnormalities in these subcortical regions are central to behavioral deficits in ASD ^40–42^. Our findings suggest that the late termination of development, or even degenerative processes in subcortical areas, may have a more profound impact on various behaviors associated with ASD than the early overgrowth observed in cortical areas. Notably, the growth trajectories of the striate cortex and neighboring regions in ASD formed inverted-U curves, reflecting ectopic abnormality. The striate cortex, located in the calcarine fissure, along with neighboring regions such as the lingual gyrus, caudal precuneus, and cuneus, is critical for processing visual information ^43^. Previous studies have indicated that increased cortical thickness in the lingual gyrus is associated with social deficits in ASD due to visual sensory abnormalities ^44^. Other studies have shown that reductions in thickness and sulcus depth of the inferior occipital cortex are associated with increased autism-related behaviors^45^. Functional magnetic resonance imaging (fMRI) responses have demonstrated that changes in receptive field properties in early visual processing areas are linked to structural changes in the occipital lobe and more severe autism symptoms ^46,47^. Nevertheless, existing reports on the inferior occipital lobe and striate cortex regions are scarce. Their extremely abnormal developmental trajectory guides us to devote more attention to the underlying neuropathological mechanisms of sensory perception abnormalities in ASD in the future.

Regional brain growth trajectories under the same category in this study may reflect different but interdependent neurodevelopmental processes. There are several possible explanations for these observations. One possibility is the relative timing of overgrowth and/or region-specific genetic mechanisms ^37^. Different brain regions may experience growth spurts at different times. This asynchronous growth can result in unique developmental trajectories for each region, even if they are functionally related. While the specific processes driving these changes remain unknown, one hypothesis is that in early development, brain systems might exhibit pluripotency, meaning they have the potential to develop into distinct network organizations. During development, neuronal populations that do not integrate into a functional domain or cognitive process are eliminated through pruning ^48^. As brain develop, they integrate within entirety while segregating from others, leading to domain specificity. This process helps refine and optimize brain networks for specific tasks and functions ^49^. Understanding the complex interplay of these factors is essential for comprehending how brain regions develop and specialize in ASD. Future research need uncover more about the specific processes driving these changes, leading to better insights into neurodevelopmental disorders.

Somewhat surprisingly, although the expected sex effect (i.e., larger brain volumes in males than in females throughout development) was observed ^50,51^, both males and females exhibited similar nonlinear trajectories across multiple global MRI phenotypes. Compared to the similar growth curves observed in global morphometric phenotypes, significant differences were found in certain cortical regions often highlighted in autism symptomatology. These areas include the inferior frontal cortex, posterior cingulate cortex, and inferior occipital lobe, which have been reported to be disturbed in autistic development. The diagnostic features of autism are similar across sexes and are likely mediated by brain regions with similar trajectories, however, the brain regions with differential developmental trajectories may explain the gender differences in symptom severity among individuals with ASD. The observed sex differences in brain morphometry may be attributed to a combination of genetic, hormonal, and environmental factors that differentially influence brain development in males and females ^25,52^. By incorporating sex-specific analyses, we achieved a more nuanced understanding of brain growth patterns, which was essential for developing tailored interventions and treatments for neurodevelopmental disorders in the future. In addition, we found significant associations between age, gender, and the degree of social deficits with individualized deviation scores from typical GMV growth curves for ASD, highlighting the spectrum nature of autism. However, given the variability in clinical assessment strategies across different sites, it remains unclear how increased brain volume is functionally important to the causal mechanisms of ASD. It is possible that abnormal brain volume is not an underlying source of core ASD symptom differences but rather a collateral consequence of the true underlying source.

ASD arises from complex interactions among various factors, encompassing both endogenous characteristics and socio-cultural aspects of the environment. Previous research has identified persistent ethnic differences in the prevalence, clinical manifestations, diagnostic criteria, education, and treatment of ASD between Western and Chinese populations ^53–55^. Although numerous studies on brain structure and function have highlighted neurological differences of TDC in China and the West ^56–58^, it remains unclear whether these differences extend to autistic brain development due to the limited availability of large Chinese ASD datasets. In this study, we constructed growth charts using overlapping age intervals from the CABIC and the ABIDE to investigate these differences. Our findings corroborate previous studies, demonstrating that Western TDCs exhibit larger brain volumes, SA, and MT compared to Chinese TDCs. Despite these differences, both groups showed similar growth trajectory shapes. Conversely, substantial variances were observed in brain morphology development among ASD individuals from different ethnic backgrounds. Ethnicity plays a critical role in shaping brain morphology in ASD. At a more resolved scale, regions exhibiting significant developmental divergences in the growth charts of ASD between CABIC and ABIDE samples are predominantly located in the cingulate gyrus and subcortical areas. These diverse abnormal developmental trajectories may reflect potential integration and divergence in brain development influenced by ethnicity.

The ABIDE samples predominantly consisted of individuals with average or above-average intelligence, whereas 70% of the autistic children in the CABIC samples exhibited intellectual disabilities. While the impact of intelligence on brain morphology during development is not fully understood, it significantly influences brain maturation ^59^. Therefore, we speculate that differences in intelligence status may contribute to the observed divergences. However, few studies have explored the moderating effects of intelligence on ASD neuroanatomy. Only some evidences suggest that individuals with Asperger’s syndrome (characterized by average or above-average IQ) exhibit milder neuroanatomical atypicality compared to those with lower IQs ^60^. In our study, we found that ethnicity, culture, and language accounted for most of the variance in WMV and SA across individuals. In contrast, for GMV, sGMV, and MT,

intelligence status appeared to be a more critical factor. Mechanistically, the processes underlying the disruption of the relationship between brain characteristics and intelligence in ASD are not yet known. This relationship may be weakened by unmeasured microstructural differences or variations in network organization. Alternatively, the factor structure of intelligence may differ in ASD, making intelligence measures less reliable compared to TDC ^61^. In conclusion, children with ASD, depending on their intellectual status, may exhibit distinct developmental trajectories, suggesting that one-size-fits-all models may not fully capture the heterogeneity of the condition. Further research is needed to explore how different levels of intellectual ability influence brain development in ASD, and to refine our understanding of these distinct trajectories. Western and Chinese children with ASD likely result from a complex interplay of factors, including race, ethnicity, intelligence characteristics, and socio-cultural environment. Future research should refine these findings, particularly examining whether the anatomical differences between children with and without ASD are causes or consequences of autism-related challenges. These parameters could potentially provide an objective basis for interpreting and diagnosing clinical symptoms of autism, necessitating extensive future cross-cultural research on the atypical growth patterns identified in this study.

### Limitation and future direction

While the CABIC provides valuable data on Chinese autistic children for replication, secondary analyses, and discovery, several limitations should be considered when applying this large-sample dataset. First, the absence of data for children under age 2.5 may have introduced bias in growth model fitting. For example, this gap could have led to exaggerated slopes at the younger age extreme in the current study. Nonetheless, our findings are supported by the results of previous longitudinal studies, which found a significantly greater rate of growth of total brain volume (TBV) between 12 and 24 months in infants at high risk of ASD, resulting in significantly greater TBV in the ASD group at age^62^. Future expansions of the CABIC dataset will try to include additional data from infants and younger children, allowing for more comprehensive coverage of early brain development in ASD.

Additionally, many sites in the CABIC dataset lacked TDC, resulting in a poor age match to ASD samples. The disparity in available TDC samples may affect the accuracy of age-matched comparisons and should be considered when interpreting findings from the CABIC dataset. While more stringent matching conditions and a case-control analysis could have been implemented, we employed normative modeling in this study to address the issue of control mismatch. This approach accounts for variability in developmental trajectories within the population, allowing for more flexible and generalized comparisons across groups. Future studies will require collaborative efforts to recruit and include a more balanced representation of TDCs to enhance the robustness and validity of cross-cultural neuroimaging research in ASD.

### Conclusion

The establishment of the CABIC marks a pivotal advancement in open collaboration for ASD research, accelerating our understanding of the neurobiological underpinnings in Chinese autistic children. The large-scale CABIC dataset will be instrumental in exploring the structural heterogeneity of ASD, offering a comprehensive understanding of the diverse neurodevelopmental patterns within the Chinese autistic population. Our preliminary analysis of this dataset addresses critical gaps in existing knowledge by focusing on early atypical brain developmental trajectories in Chinese autistic children and providing a novel cross-cultural perspective on ASD. The insights gained from this extensive dataset will support the development of precise clinical support and educational strategies tailored to the unique needs of autistic children, ultimately enhancing their quality of life.

## Methods

### Contributing Sites

CABIC was initiated in 2023 and remains open to new members, with the goal of aggregating existing MRI data from a wide range of autism research groups in China. The consortium initially included 16 sites to establish collaboration, and 11 sites had approval to share individual data. Written informed consent from participants or their guardians was obtained, with approval from the local ethics committees for each dataset. Deidentified and anonymized data were contributed from studies approved by local Institutional Review Boards. All contributing sites had approval to share anonymized individual data.

### Phenotypic Data

Prior to data aggregation, consortium members agreed on a basic phenotypic protocol by identifying overlapping measures across sites. These measures included age at scan, sex, intelligence scores (which was estimated by the Chinese Version of Wechsler Intelligence Scale, development quotient or Peabody Picture Vocabulary Test 4th Edition), handedness, and diagnostic information. Basic demographic and clinical information for the cohort is summarized in **Table S1** and **Figure 1**. Contributors were encouraged to provide as many additional measures as possible, although such contributions were not required due to the voluntary, unfunded nature of this effort. Inclusion as an autistic participant required a diagnosis of Autistic Disorder by an experienced clinician, based on DSM-IV or DSM-5 criteria, supplemented by the CARS, the ADOS, the ADI-R, ABC, SRS and/or RBS-R. Children with ASD-related medical conditions (e.g., Fragile X syndrome, tuberous sclerosis) or other neurological conditions (e.g., epilepsy, Tourette’s syndrome) were excluded. See **Supplementary Information** for further information on participants and assessments from all sites.

### Structural MR Imaging data

Structural T1-weighted brain MRI scans were collected at each study site using a variety of scanners and protocols at field strengths of 3 T. During MRI scanning, all autistic children from all sites were sedated with 50 mg/kg chloral hydrate. The TDCs from six of the eleven sites (UESTC, PLAGH, SYSU, JNU, YZU, BNU) were instructed to lie still and watch cartoons or keep their eyes closed to rest. The TDC from the other five sites (CHCMU, HMU, PKU, JLU, GMCMC) were given chloral hydrate sedation. Sedation was performed by a trained and certified nurse following the guidelines and protocols established by the Radiology Sedation Committee of the local hospital. A caregiver was present for each participant during the scan.

### Image data processing pipeline

#### Step 1: Segmentation

The MRI scans were segmented using a standardized automated pipeline based on FreeSurfer (version 6.0) to obtain general morphological measurements of different brain tissues from *aseg.stats* and *lh/rh.aparc.stats*. These morphometric phenotypes included total GMV, sGMV, WMV, mean CT, and SA. Regional neurotypes including GMV were also obtained according to the Desikon-Killiany (DK) and *Aseg* parcellation. To estimate the overall developmental charts of the brain region in subsequent analyses, we averaged the symmetrical brain regions of the right and left cerebral hemispheres.

#### Step 2: Image quality control

Firstly, T1 images were visually inspected for quality by two experienced raters, and participants with lower-quality data were excluded. All the images passed this quality control. Secondly, any scan that could not undergo the entire data processing pipeline was excluded, resulting in the removal of 66 participants. Finally, the Euler number was utilized to assess the quality of the reconstructed cortical surface. The Euler number is a mathematical concept that summarizes the topological complexity of a surface and can be calculated as 2-2n, where n represents the number of defects such as holes or handles. A high Euler number represents a surface with fewer defects, indicating high-quality cortical surface reconstruction. The Euler number is a reliable and quantitative measure and can be used to identify unusable images ^7,63^.

#### Step 3: Multisite data harmonization

In studies involving multisite datasets, it is critical to account for and remove site-specific effects to estimate generalizable results from multi-site or multi-study neuroimaging data. In recent years, the *ComBat* method and its variants have been increasingly used to achieve harmonization of MRI data acquired across multiple sites^64^. In the current study, ComBat with a generalized additive model (GAM) was employed for harmonization between different studies (Some sites include multi-studies data obtained from different scanners, check the supplementary information for details). This approach allowed us to remove study-specific differences in individual morphometric phenotypes while retaining the ability to conduct downstream nonlinear modeling analyses ^65^. Age, diagnostic group and sex were included as biological variables that needed to be protected, with age set as a smooth term.

### Modeling normative trajectories

By using the *gamlss* package (version 5.4-3) in R 4.2.0, GAMLSS was adopted to estimate the age-dependent brain morphological growth charts.

#### Model data distributions

While the World Health Organization provides guidelines for anthropometric growth chart modeling with GAMLSS, the LBCC recently reported that the generalized gamma distribution offered the best fit for brain morphometry measures ^7^. Consequently, the same GAMLSS with generalized gamma distributions was applied for the trajectories in the current study across all evaluated models.

#### Fractional polynomial model set

We constructed the GAMLSS using the code that has been generated by LBCC (https://github.com/brainchart/lifespan). This modified sex-stratified GAMLSS approach allowed modeling of age-related changes in brain morphometry measures while considering the random effects of sites. The random effects are drawn from a normal distribution with zero mean and variance to be estimated, *γ ∼Ν (0, δ*^2^). The ability to include random effects is fundamental to accounting for co-dependence between observations. Additionally, the generalized gamma distribution, which has four parameters: median (*µ*), coefficient of variation (*σ*), skewness (𝜈), and kurtosis (*τ*), was chosen to accommodate the data distribution. The GAMLSS framework of each brain morphometry measure, denoted with Y, could be specified in the following way:

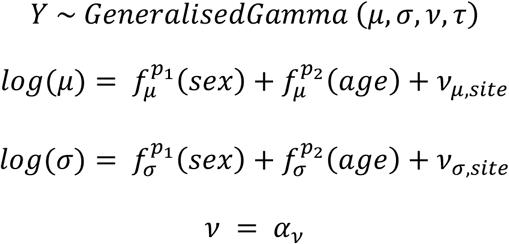

Each fractional polynomial, *f(),* can be viewed as a simpler form of spline modelling using a fixed set of polynomials. The terms “order”, *p*, is used to refer to the number of terms in the fractional polynomial model. The GAMLSS framework includes the ability to estimate the most appropriate powers of fractional polynomial expansion within the iterative fitting algorithm, searching across the standard set of powers, *p*∈ (−2, −1, −0.5, 0, 0.5, 1, 2, 3). As conventionally defined by previous study ^66^, a second order fractional polynomial where power *p* is repeated is defined as *β*_*p*_*x*^*p*^ + *β*_*p*′_*x*^*p*^ *log*(*x*), while a third order fractional polynomial where power *p* is repeated is defined as *β*_*p*_*x*^*p*^ + *β*_*p*’_*x*^*p*^ *log*(*x*) + *β*_*p*′_*x*^*p*^ *log* (*x*)^2^. No smoothing terms were used in any GAMLSS models implemented in this study

For Model convergence within GAMLSS, we used the default convergence criterion of log-likelihood = 0.001 between model iterations. Due to the limitations of the traditional bootstrapping procedure, it is possible for a bootstrap replication to become degenerate, meaning the resampled subset of data causes model fitting instability or non-convergence. Therefore, we followed the method of Bethlehem et al., employing a stratified bootstrap procedure ^7^. We used the Bayesian Information Criterion (BIC) to compare model fits among the converged models, with a lower BIC indicating a better fit. The optimal model for a given brain morphometry measure was selected based on the lower BIC value among all convergent models. In addition to this, given that LBCC has previously mapped standard brain growth across the lifespan using a much larger sample of healthy participants, we ultimately chose to use the same parameters for our model estimation as they did. This decision was made to ensure consistency and comparability in our subsequent validation efforts. By aligning our parameters with those used by LBCC, we aimed to facilitate a direct comparison between our results and their established brain growth maps. This approach not only enhances the robustness of our findings but also situates our work within the broader context of existing research in the field.

Across all global morphometric phenotypes (GMV, sGMV, WMV, SA, MT), modeling indicated 3rd order fractional polynomial fits for the *µ*-component, and 2nd order fractional polynomial fits for the *σ*-component. In addition, Bethlehem et al. indicated that fractional polynomial modeling for 𝜈 resulted in model instability. Therefore, we also adopted the inclusion of an intercept term only for the 𝜈 component for all brain morphometry measures. Then, we generalized the same GAMLSS modeling approach to estimate trajectories for regional volume at each of 34 cortical areas (all brain regions in *DK* parcellation) and 12 subcortical areas (including thalamus, caudate, putamen, pallidum, hippocampus, amygdala and different regions of cingulate cortex). To visualize the developmental changes in local brain regions, we mapped relative volume values on the cortical surface. Initially, we used existing developmental samples of Chinese children and adolescents from the Chinese Color Nest Project to determine the peaks of developmental volume values^67,68^. For each child in our study, we calculated the difference in volume values for each brain region compared to their peak values in corresponding brain regions, defining these as relative volume values. We then divided both TDC and autistic children into five age groups and averaged the relative volume values within these groups.

### Defining developmental shift milestones

Previous research has led to the theory of age-specific anatomical abnormalities in ASD, demonstrating abnormal brain overgrowth during early ages, followed by an accelerated rate of decline and possible degeneration in later ages in this condition. GAMLSS modeling enabled us to leverage the large neuroimaging dataset to identify this developmental shift milestone. This milestone indicated the turning point from over to delayed maturation in autistic children compared to typically developing children (i.e., the intersection of two curves). Key points were determined based on the GAMLSS model output (50th centile) for each of the global morphometric phenotypes. To analysis trajectories and milestones of brain development with finer-grained anatomical resolution, a similar series of methodological steps was performed for the regional morphometric phenotype.

Additionally, a shape-based whole time-series clustering algorithm was used to identify different categories of abnormal developmental trajectories across all brain regions ^21^. Firstly, we calculated the regional difference curves between the ASD and typically developing children’s development trajectories. Then, the *k*-Shape algorithm was used on regional difference curves to cluster all brain regions into different abnormal developmental categories ^69^.

### Individualized deviation centile scores and inter-subject similarity analysis

In addition to constructing developmental curves for ASD and TDC separately, quantified each autistic children’s relative distance from reference models of typical development are important for understanding ASD. After establishing normative reference ranges using the GAMLSS model in TDC, individual deviation centile scores of autistic children were estimated in terms of its relative distance from the median of the age-normed and sex-stratified distributions provided by the reference model. Individually deviation centile scores have been used by the LBCC to investigate case-control differences across disorders ^7^ (See supplementary method in *Bethlehem et.al, 2022* for further details). This approach was conceptually similar to the previously reported quantile rank mapping, with atypical individual having more extreme percentile scores (or quantiles) ^15,70^.

RSA was used to test whether the individual deviation centile scores correlate with individual variation in the autistic children’s behavioral and clinical characteristics ^71,72^. Firstly, we calculated the pairwise similarity between participants with respect to the individual deviation centile scores by Pearson correlation, and obtained a centile scores similarity matrix. Then, this process was repeated for different demographic variables and behavioral measurements, including age, sex, social deficits scores (measured by social responsiveness scale), scanning-site. Notably, the modelling procedure differed depending on the data-scale (categorical- or dimensional-valued data). For dimensional-valued measurements, such as age and social deficits scores, subject distance was the absolute Euclidean distance between their measurements. For categorical variables (such as sex or scanning-site) subject distance was 0 if categorical variables matched (same sex or same scanning-site) and 1 otherwise. Finally, Kendall rank correlation coefficient (Kendall *τ*) was used to compared the centile scores similarity matrix to the matrices for each individual characteristic. To construct RSA models, we used data from all autistic children that had the scores on that measurement.

To investigate the relationship between regional individual deviations in specific brain regions and SRS scores, we employed the *Glmnet* package in R (https://glmnet.stanford.edu/articles/glmnet.html), a toolkit designed for fitting generalized linear models via penalized maximum likelihood. All individual deviation scores were used as predictor variables in a penalized regression model, with SRS scores as the outcome variable. By utilizing the elastic net regularization provided by *Glmnet*, we traversed a range of lambda values to optimize the model, balancing the trade-off between model complexity and predictive accuracy. Each model was created using a 10-folds cross-validation with the cv.glmnet function from glmnet package. This approach allowed us to identify the most significant brain regions while excluding non-contributory ones, resulting in a sparse model where only a few significantly contributed brain regions were retained as significant predictors of age, sex or SRS scores.

### Replication analysis

To illustrate the autistic growth curve differences between populations, ABIDE samples between 5-18 years were adopted to constructed the growth curves by GAMLSS model. Due to the limitation of relatively smaller sample size of intellective subgroups, MFPR was used to generate optimized intelligence-specific models. This algorithm was implemented using the “*mfp*” package in R. Recent study has indicated that MFPR had optimal performance for generating normative trajectory modeling when analyzing small datasets ^73^.

To quantitatively estimate the diversity in brain growth attributable to ethnicity (referring between CABIC to ABIDE), we computed the NV of global morphometric phenotypes and regional GMV with the following equation:

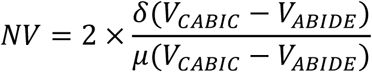

where *V* is a vector referring to the morphometry measure and *δ* refers to the standard deviation. NV indicates the degree of curve shape dispersion between two growth curves across ages (sampling interval = 0.1). *δ* is normalized by the mean volume of parcels across two samples, denoted as *μ*. A small NV indicates that two brain growth curves share similar shapes, and vice versa ^15,68^.

## Data availability

All datasets from the CABIC will be made publicly available through the national scientific database in the future (https://www.scidb.cn/en). At this stage, access to the data is restricted to researchers and collaborators within CABIC. However, any institution or researcher can request access to the full dataset across all participating sites by applying for CABIC membership (https://php.bdnilab.com/resources/#;). CABIC actively encourages broader participation and welcomes more institutions and research sites to join this collaborative effort, fostering a more comprehensive and inclusive resource for advancing autism research.

## Code availability

All code is available at https://github.com/CABIC-COM/Charting-Brain-Growth-in-Chinese-Children-with-Autism.

## Acknowledgements

X.D. was supported by National Key R&D Program of China(2024YFF0509100), National Natural Science Foundation of China (82322035, 62273076), Key-Area Research and Development Program of Guangdong Province (2023B0303020001). X.Z. was supported by the STI 2030 - the major projects of the Brain Science and Brain-Inspired Intelligence Technology (2021ZD0200500), the Key-Area Research and Development Program of Guangdong Province (2019B030335001). H.C. was supported by National Natural Science Foundation of China (62333003, 62036003). T.L. was supported by Guangdong Key Project (2018B030335001). J.C. was supported by the Special Key project of Technology Innovation and Application Development of Chongqing Science and Technology Bureau (No. CSTB2022TIAD-KPX0151). L.W. and J.W. were supported by National Natural Science Foundation of China (U24A20770, 81874270). F.J. was supported by Key R&D Project of the Jilin Provincial Department of Science and Technology (20230203067SF), Innovation Team Project under the Talent Special Program of the Jilin Provincial Department of Science and Technology (20240601003RC). A.C. was supported by Key Programs of the National Social Science Foundation. S.L. was supported by National Key R&D Program of China (2022YFC2409404), Crossing Projects of Beijing Nova program (20240484674), and Capital’s Funds for Health Improvement and Research (CFH2024-2-5024) and National Natural Science Foundation of China (81971603). X.Y. was supported by the “Science and Technology Innovation 2030” - “Brain Science and brain-like Research” major project from the Chinese Ministry of Science and Technology (2021ZD0200522), and Guangzhou Science and Technology Plan Project (2023A03J0900). X.W. was supported by National Natural Science Foundation of China (52400247). J.J. was supported by Guangdong Basic and Applied Basic Research Foundation (2022B1515130007). R.Z. was supported by grants from Beijing Natural Science Foundation (J230013). H.J. was supported by the Wuxi Municipal Health Commission’s ‘Double Hundred’ Medical Health Young Elite Talent Project (Grant No. BJ2023088).

## Author contributions

X.D. initiated the formation of the CABIC. L.L and X.D. designed the study. L.L conducted analyses and wrote intimate manuscript. X.D. and X.Z. reviewed and edited manuscript. All other authors made substantial contributions to the conception or design of the work, the acquisition, analysis or interpretation of data, or drafted or substantively revised the Article.

## Supplementary Methods

### Sensitivity analyses of GAMLSS

The resulting models were evaluated using several sensitivity analyses and validation approaches:

1. **Leave-one-site-out:** We performed a series of leave-one-site-out analyses to test whether our model’s reliability was skewed toward any study. Specifically, we iteratively excluded the dataset from a single site (e.g., UESTC) from the primary studies, refitted the GAMLSS model, re-estimated all model parameters, and then extracted developmental trajectories. These alternative trajectories were then compared to those derived from the full dataset for all brain morphometry measures. Our results showed remarkable consistency (**Figure S1**), with a high degree of similarity between the trajectories derived from the full dataset and those from the subsets.
2. **Bootstrap analysis:** To further assess the reliability of the GAMLSS-fitted trajectories and obtain their confidence intervals, we conducted a bootstrap resampling analysis. Specifically, we performed 1,000 bootstrap iterations with stratified sampling and replacement. The bootstrap replicates were stratified by site and sex, preserving the relative proportions of the original datasets. For each brain morphometry measure, we refitted 1,000 trajectories and computed 95% confidence intervals (CIs) for the 50th centile curve. Supplementary **Figure S2** depicts the CIs for the developmental trajectories, underscoring the stability of our modeling framework.
3. **Adjusted Lifespan Brain Chart Consortium (LBCC) brain chart:** Based on the lifespan trajectory from the LBCC’s seminal work, an online tool has been developed for the lifespan development of human brain morphology (https://brainchart.shinyapps.io/brainchart/). ASD and TDC samples were employed separately to update the LBCC charts.
4. **Longitudinal variability:** Due to the lack of longitudinal imaging data, normative models were estimated from cross-sectional data collected at a single time point. However, with the available longitudinal data from 28 TDCs from BJN (three follow-ups) and 17 autistic children from YZU (two follow-ups), we further tested the generalizability of different models to capture within-subject variation in centile scores over time, quantified as the interquartile range (IQR). Our findings revealed that the IQR for centile scores was consistently low (all median IQR < 0.05 centile points), indicating highly stable centile scoring across multiple repeated scans. In addition to the possibility of constructing own brain growth curves using our large-sample data, the LBCC provided an online tool (https://github.com/brainchart/lifespan) for uploading local data to update and enhance the LBCC normative brain growth charts. Here, we employed CABIC samples into this procedure to obtain an adjusted LBCC brain growth chart. We found that across all the morphometric phenotypes, the IQR of our CABIC brain growth chart was significantly decreased compared to the IQR of the upgraded LBCC brain growth chart (Two-sample T-test, *P* < 0.05) (**Figure S3**). These results indicated that our CABIC-generated brain growth charts show reliable stability and align more closely with the growth and developmental trajectories of Chinese children.

### Quality control

Initially, T1 images were visually inspected for quality by two experienced raters, and participants with lower-quality data were excluded. All remaining images passed this quality control process. Subsequently, any scan that could not be fully processed through the data pipeline was excluded, resulting in the removal of 66 participants. Finally, we assessed the impact of image quality on estimated brain phenotypes and the parameterization of the GAMLSS model using the Euler Index (EI), an automated and quantitative measure of data quality in scans processed by FreeSurfer.

The EI metric was defined as the sum across hemispheres of the number of surface “holes” or topological defects in the cortical surface reconstruction before the topological correction performed by the FreeSurfer pipeline. Significant differences in EI were observed across sites (**Figure S4a**, ANOVA, *p* < 0.0001) and between ASD and TDC groups (**Figure S4b**, two-sample *t*-test, *p* < 0.0001). We also examined the relationship between image quality, as measured by EI, and individual deviation centile scores of GMV using Spearman correlation analysis. No significant associations were found between EI and individual centile scores (**Figure S4c**). Notably, although there were significant between-group differences between ASD and TDC groups within CABIC, no significant differences were observed between the two large databases, ABIDE and CABIC (**Figure S4d**, ANOVA, *p* = 0.65).

### Sexual variability in trajectories

In the GAMLSS modeling process, the sex effect was incorporated as a critical feature to establish sex-stratified growth curves for children. Male predominance is a well-documented characteristic of ASD, with a reported male-to-female ratio of approximately 4:1. The data distribution in our study similarly exhibited a male preponderance among autistic children. Consequently, the primary results focus on the growth curves for males, which are presented in the main text, while those for females are provided in Supplementary **Figures S7-S8**. Furthermore, the regional sex-stratified growth curves for ASD are illustrated in **Figure S9**, while for TDC are illustrated in **Figure S10**, offering valuable insights into the differential effects of sex on ASD development.

### Intelligence groupings

To characterize intelligence-specific growth charts, we categorized all autistic children in the CABIC dataset into subgroups based on intelligence quotient (IQ) cutoff scores. It is important to note that IQ assessment methods vary across different sites, leading to potential inconsistencies. An IQ assessment was considered valid if it employed one of the following standardized measures: the Wechsler Intelligence Scale for Children (WISC), Gesell Developmental Schedules (GDS), or the Peabody Picture Vocabulary Test (PPVT).

To standardize groupings across these different scales, specific cutoff scores were applied. For the WISC, children with a Full-Scale Intelligence Quotient (FIQ) score below 90 were classified into the intellectual disability group, while those with scores of 90 or above were classified into the average intelligence group; For the GDS, children with a Developmental Quotient (DQ) score below 85 were placed in the intellectual disability group, while those with scores of 85 or above were classified into the average intelligence group; For the PPVT, children with standardized scores below 90 were categorized into the intellectual disability group, and those with scores of 90 or above were classified into the average intelligence group.

### Reference multi-sites database details

Site-specific details for further information on ethics, diagnostics and assessments. are available at https://php.bdnilab.com/sites/. The detailed acquisition parameters of each site are presented following:

### UESTC

#### STUDY1: 3.0-T GE Discovery MR750 scanner

High-resolution MR data of the whole brain were acquired using a 3D T1 sequence with the following parameters: repetition time (TR) = 6.02 ms, echo time (TE) = 1.96 ms, matrix size = 256 x 256, flip angle (FA) = 9°, field of view (FOV) = 256 x 256 mm^2^, voxel size = 1 x 1 x 1 mm^3^, and 156 axial slices.

#### STUDY2: 3.0-T Siemens Skyra MRI scanner

High-resolution MR data of the whole brain were acquired using a 3DT1 sequence with the following parameters: TR = 1570ms, TE = 2.4ms; matrix size = 256 x 256; FA = 8°; FOV = 256 x 256 mm^2^; voxel size = 1 x 1 x 1 mm^3^, slice number=160.

#### STUDY3: 3.0-T Siemens Skyra MRI scanner

High-resolution MR data of the whole brain were acquired using a 3DT1 sequence with the following parameters: TR = 1570ms, TE = 2.4ms; matrix size = 256 x 256; FA = 8°; FOV = 256 x 256 mm^2^; voxel size = 1 x 1 x 1 mm^3^, slice number=160.

#### STUDY4: 3.0-T UNITED IMAGING uMR780

High-resolution MR data of the whole brain were acquired using a 3DT1 sequence with the following parameters: TR = 6.9ms, TE = 3ms; matrix size = 256 x 256; FA = 10°; FOV = 256 x 256 mm^2^; voxel size = 1 x 1 x 1 mm^3^, slice number=176.

### SYSU

#### 3.0-T Siemens Skyra MRI scanner

High-resolution MR data of the whole brain were acquired using a 3DT1 sequence with the following parameters: TR = 1800ms, TE = 2.19ms; matrix size = 256 x 256; FA = 9°; FOV = 256 x 256 mm^2^; voxel size = 1 x 1 x 1 mm^3^, slice number=176.

### WXCH

#### Siemens Magnetom Aera scanner

High-resolution MR data of the whole brain were acquired using a 3D T1W1 sequence with the following parameters: TR = 2200ms, TE = 3.06ms; FOV = 192 x 192 mm^2^; slice thickness =1mm.

### PKU

#### 3.0-T GE Discovery MR750 scanner

High-resolution MR data of the whole brain were acquired using Ax FSPGER BRAVO sequence with the following parameters: TR = 8.2ms, TE = 3.2ms; matrix size = 256 x 256; FA = 9°; FOV = 256 x 256 mm^2^; voxel size = 1 x 1 x 1 mm^3^, slice number=192, prep time = 600.

### HMU

#### 3.0-T Philips Achieva MRI scanner

High-resolution MR data of the whole brain were acquired using a 3DT1 sequence with the following parameters: TR = 8.4ms, TE = 3.87ms; matrix size = 256 x 256; FA = 9°; FOV = 256 x 256 mm^2^; slice thickness = 1mm, voxel size = 1 x 1 x 1 mm^3^, slice number=176.

### YZU

#### 3.0-T GE Discovery MR750 scanner

High-resolution MR data of the whole brain were acquired using a 3DT1 sequence with the following parameters: TR = 7.2ms, TE = 3.06ms; matrix size = 256 x 256; FA = 12°; FOV = 256 x 256 mm^2^; slice thickness = 1mm, voxel size = 1 x 1 x 1 mm^3^, slice number=166.

### JLU

#### 3.0-T Philips Ingenia Elition scanner

High-resolution MR data of the whole brain were acquired using a 3DT1 sequence with the following parameters: TR = 6.7ms, TE = 3.06ms; matrix size = 240 x 240; FA = 8°; FOV = 240 x 240 mm^2^; slice thickness = 1mm, voxel size = 1 x 1 x 1 mm^3^, slice number=152.

### CHCMU

#### 3.0-T Philips Achieva MRI scanner

High-resolution MR data of the whole brain were acquired using a 3DT1 sequence with the following parameters: TR = 7.7ms, TE = 3.8ms; matrix size = 256 x 256; FA = 12°; FOV = 256 x 256 mm^2^; slice thickness = 1mm, voxel size = 1 x 1 x 1 mm^3^, slice number=164.

#### 3.0-T Siemens Trio MRI scanner

High-resolution MR data of the whole brain were acquired using a 3DT1 sequence with the following parameters: TR = 8.22ms, TE = 3.19ms; matrix size = 256 x 256; FA = 12°; FOV = 256 x 256 mm^2^; slice thickness = 1mm, voxel size = 1 x 1 x 1 mm^3^, slice number=164.

### PLAGH

#### 3.0-T GE Discovery MR750 scanner

High-resolution MR data of the whole brain were acquired using a 3DT1 sequence with the following parameters: TR = 7.9ms, TE = MIN FULL; matrix size = 256 x 256; FA = 12°; FOV = 256 x 256 mm^2^; slice thickness = 1mm, voxel size = 1 x 1 x 1 mm^3^, slice number=156.

### CMCMC

#### 3.0-T Philips Healthcare scanner

High-resolution MR data of the whole brain were acquired using a 3DT1 sequence with the following parameters: TR = 8.1ms, TE = 3,7ms; matrix size = 256 x 240; FA = 8°; FOV = 256 x 240 mm^2^; slice thickness = 1mm, voxel size = 1 x 1 x 1 mm^3^, slice number=160.

### BNU

#### STUDY1:3.0-T Siemens Trio MRI scanner

High-resolution MR data of the whole brain were acquired using 3D MPRAGE sequence with the following parameters: TR = 2600 ms, TE = 3.02ms; matrix size = 256 x 256; FA = 8°; FOV = 256 x 256 mm^2^; slice thickness = 1mm, voxel size = 1 x 1 x 1 mm^3^, slice number=176.

#### STUDY2:3.0-T GE Discovery MR750 scanner

High-resolution MR data of the whole brain were acquired using 3D SPGR sequence with the following parameters: TR = 6.7 ms, TE = 2.9 ms; matrix size = 256 x 256; FA = 12°; FOV = 256 x 256 mm2; slice thickness = 1mm, voxel size = 1 x 1 x 1 mm3, slice number=176.

## Supplementary Figures

**Figure S1.**
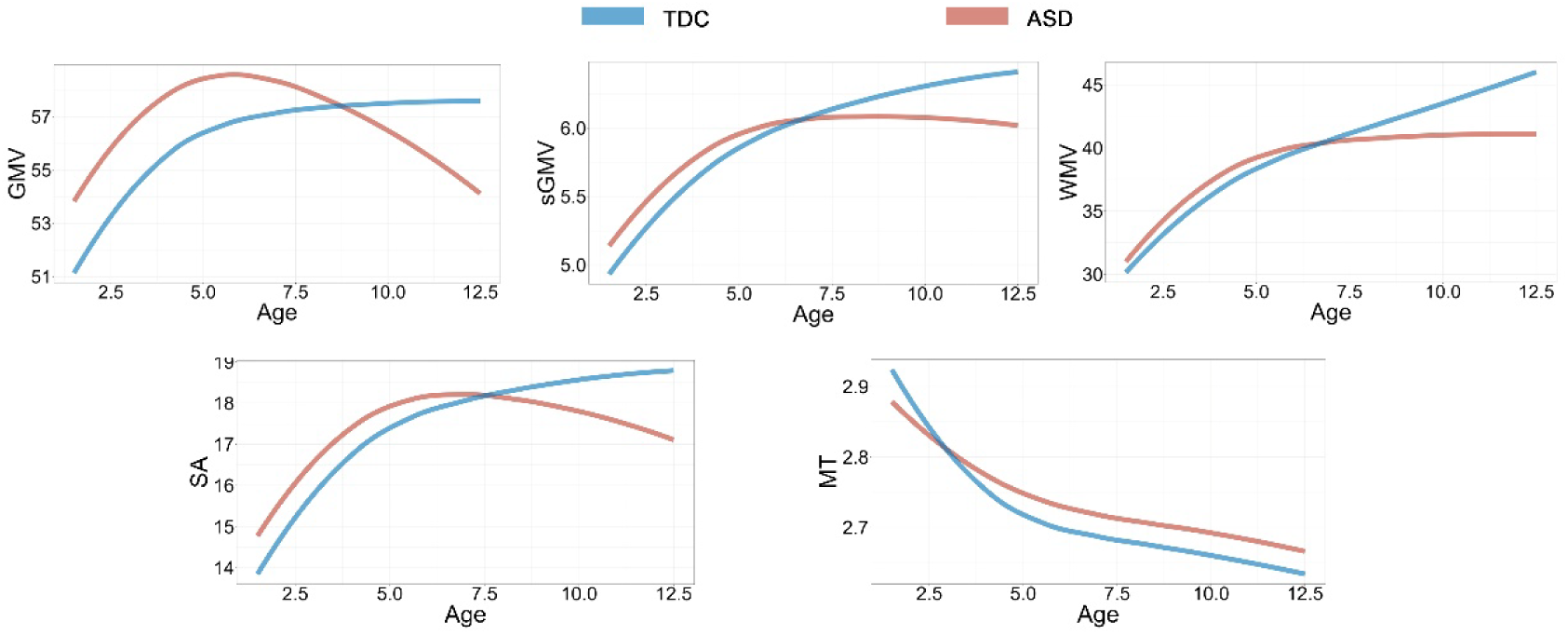
Leave-one-site-out (LOSO) analyses of typical and atypical trajectories in males for global morphometric phenotypes. The solid lines represent the 95% confidence intervals, computed from the mean and standard deviation of normative trajectories, repeatedly estimated after systematically excluding data from each contributing site in turn.

**Figure S2.**
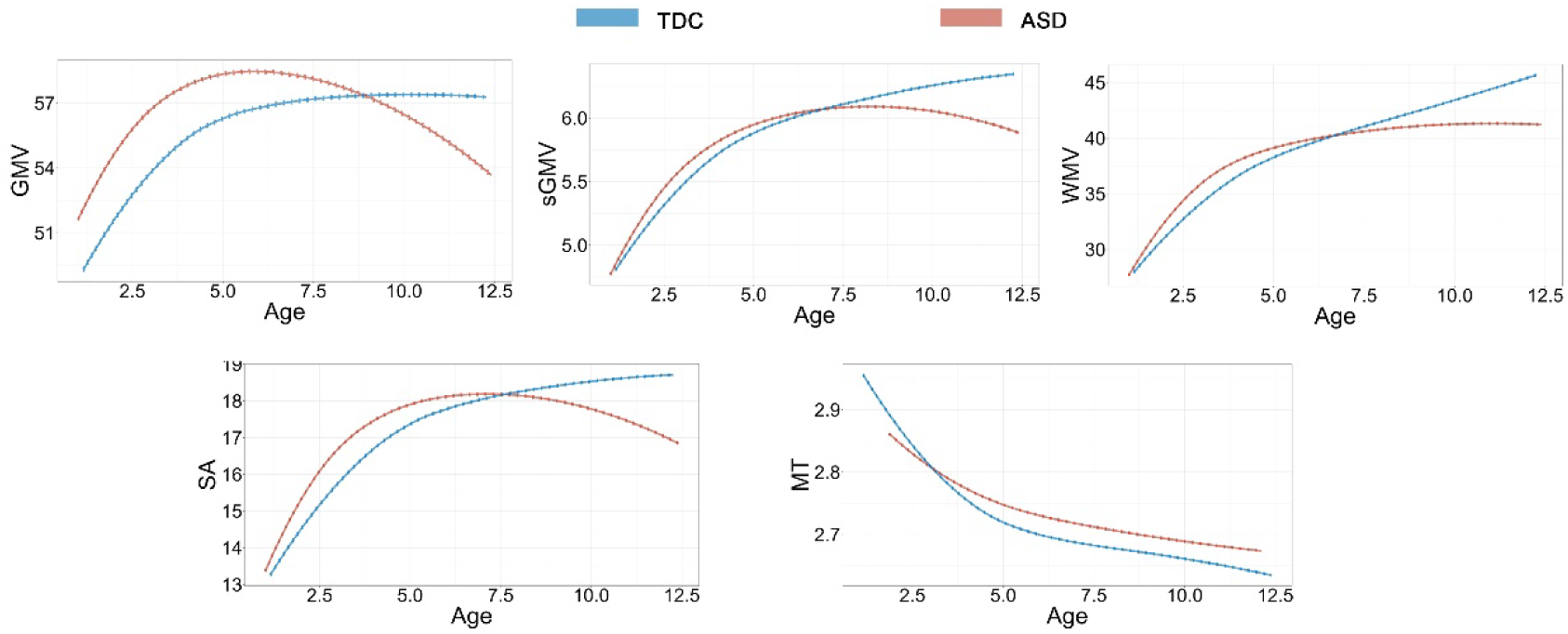
Bootstrap resampling of confidence intervals on typical and atypical trajectories in males for global morphometric phenotypes. The median (50% centile) is represented by a solid line, while the 95% centiles are indicated by dotted lines. 95% confidence intervals (estimated across random bootstrap iterations resampling with replacement) were computed from the mean and standard deviation of trajectories after 1000 iterations of a bootstrapping procedure designed to conserve the relative proportion of primary sites in each resampling with replacement from the representative dataset.

**Figure S3.**
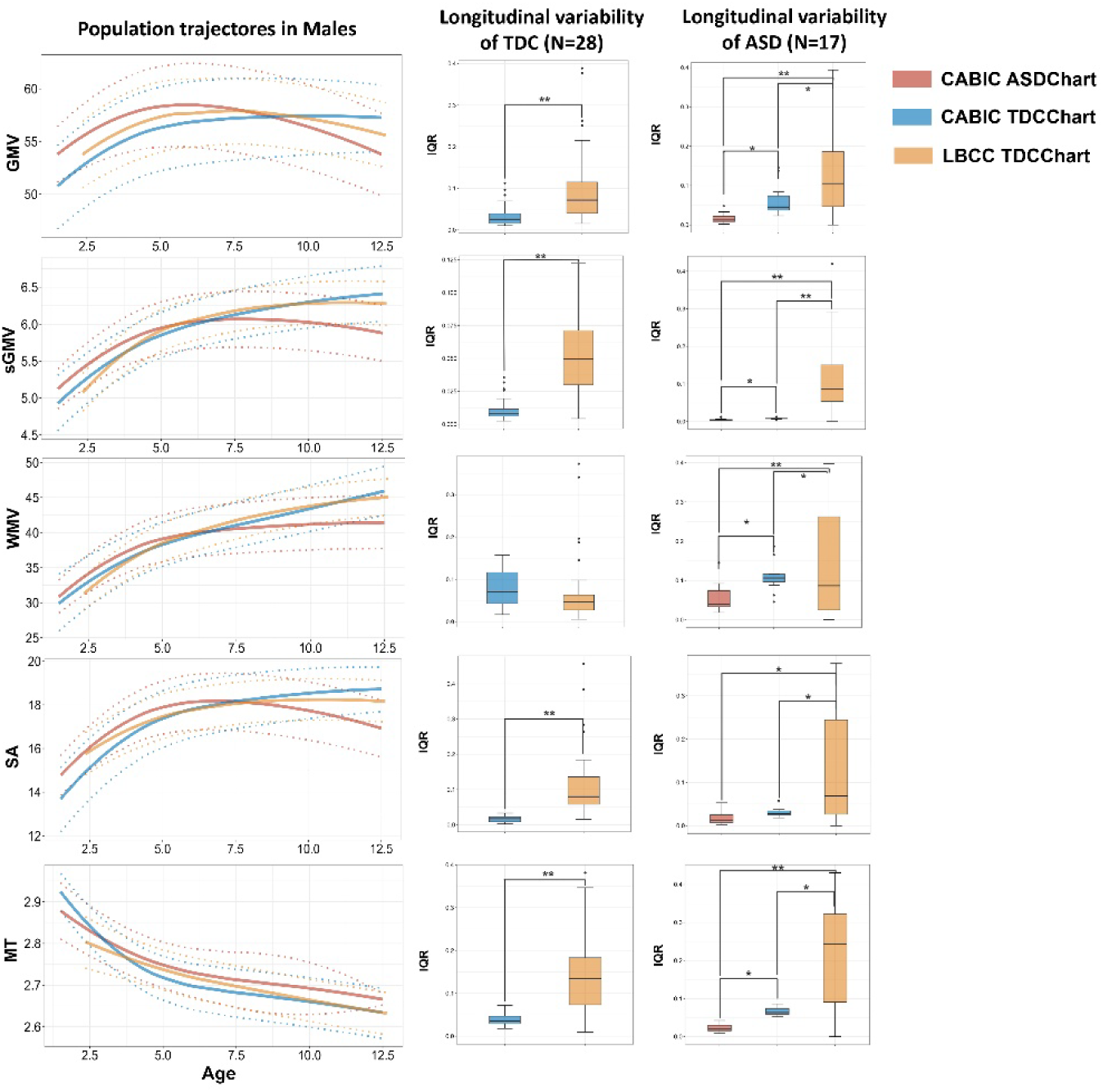
Comparison of brain charts independently estimated from the CABIC dataset with previously published brain charts from the Lifespan Brain Chart Consortium (LBCC) Left: Trajectories of global morphometric phenotypes. Right: Within-subject variation, quantified as the interquartile range (IQR) in longitudinal centile scores for global morphometric phenotypes, comparing different brain chart curves.

**Figure S4.**
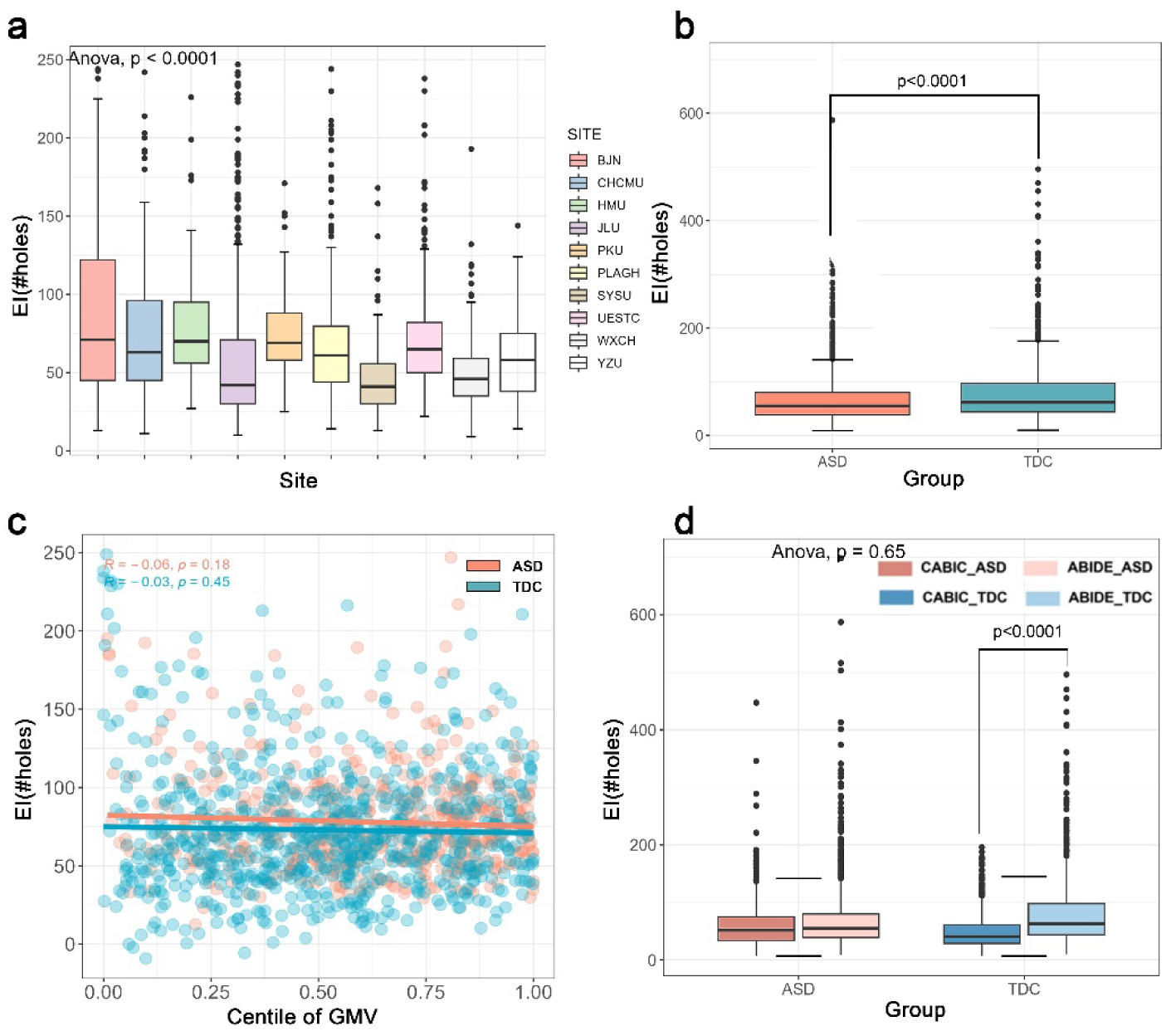
Complementary analyses of quality control metrics defined by the Euler Index (EI) **(a)** Bar plots showing the distribution of EI across all sites. **(b)** Significant between-group differences in EI between ASD and TDC groups. **(c)** Associations between individual deviation centile scores of GMV and MRI scan quality, as defined by EI. The Spearman correlations between EI and individual centile scores of GMV were negligible (ASD, *p* = 0.18; TDC, *p* = 0.45). **(d)** Differences in EI between the ABIDE and CABIC databases.

**Figure S5.**
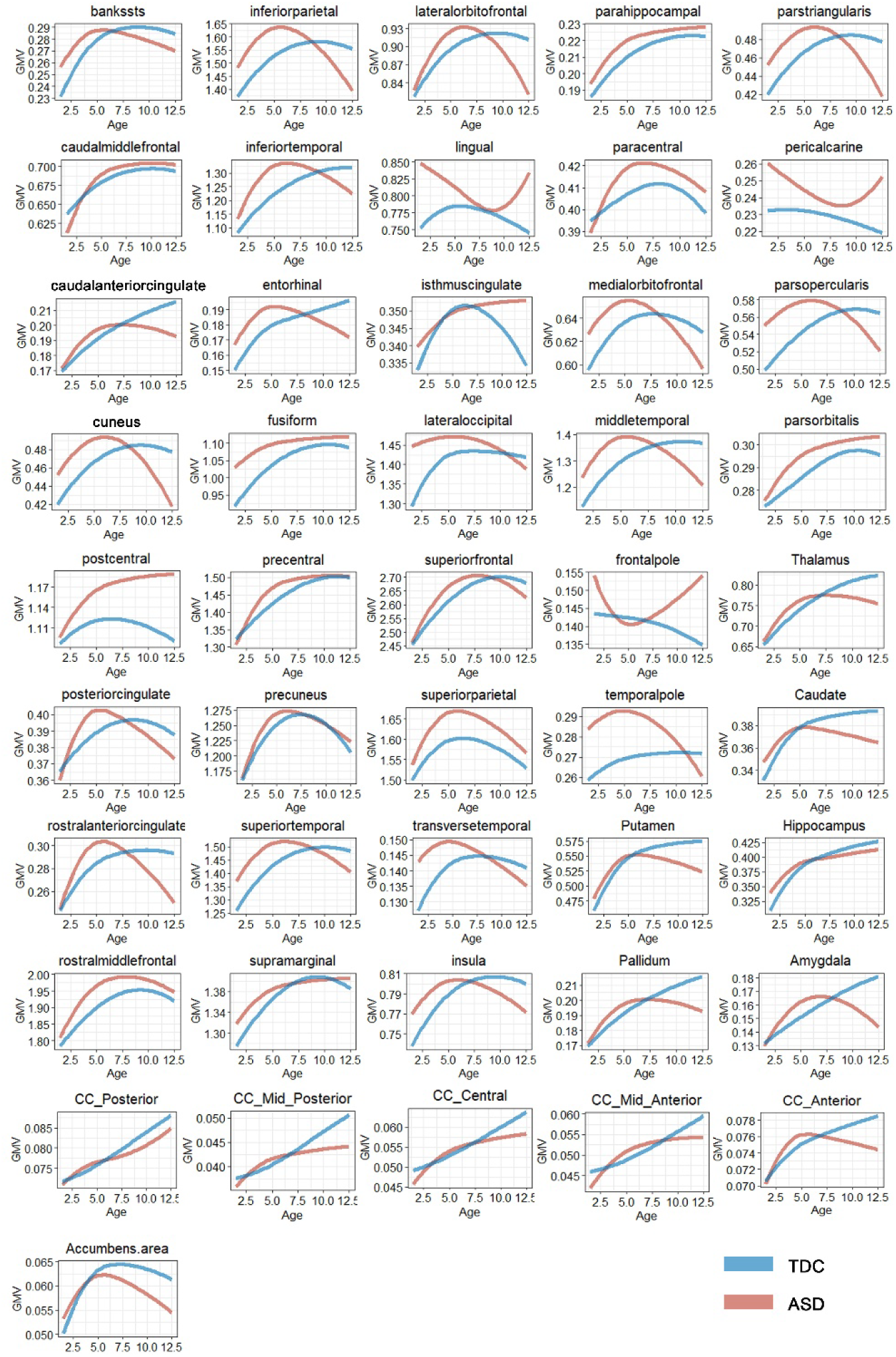
Typical and atypical trajectories of median regional volumes in males for 34 bilateral brain regions as defined by the Desikan-Killiany parcellation and 12 subcortical regions (with volumes normalized to 10,000 mm³) These trajectories were fitted to the raw data using the same GAMLSS model applied to estimate the global morphometric phenotype trajectories, as illustrated in Figure 2.

**Figure S6.**
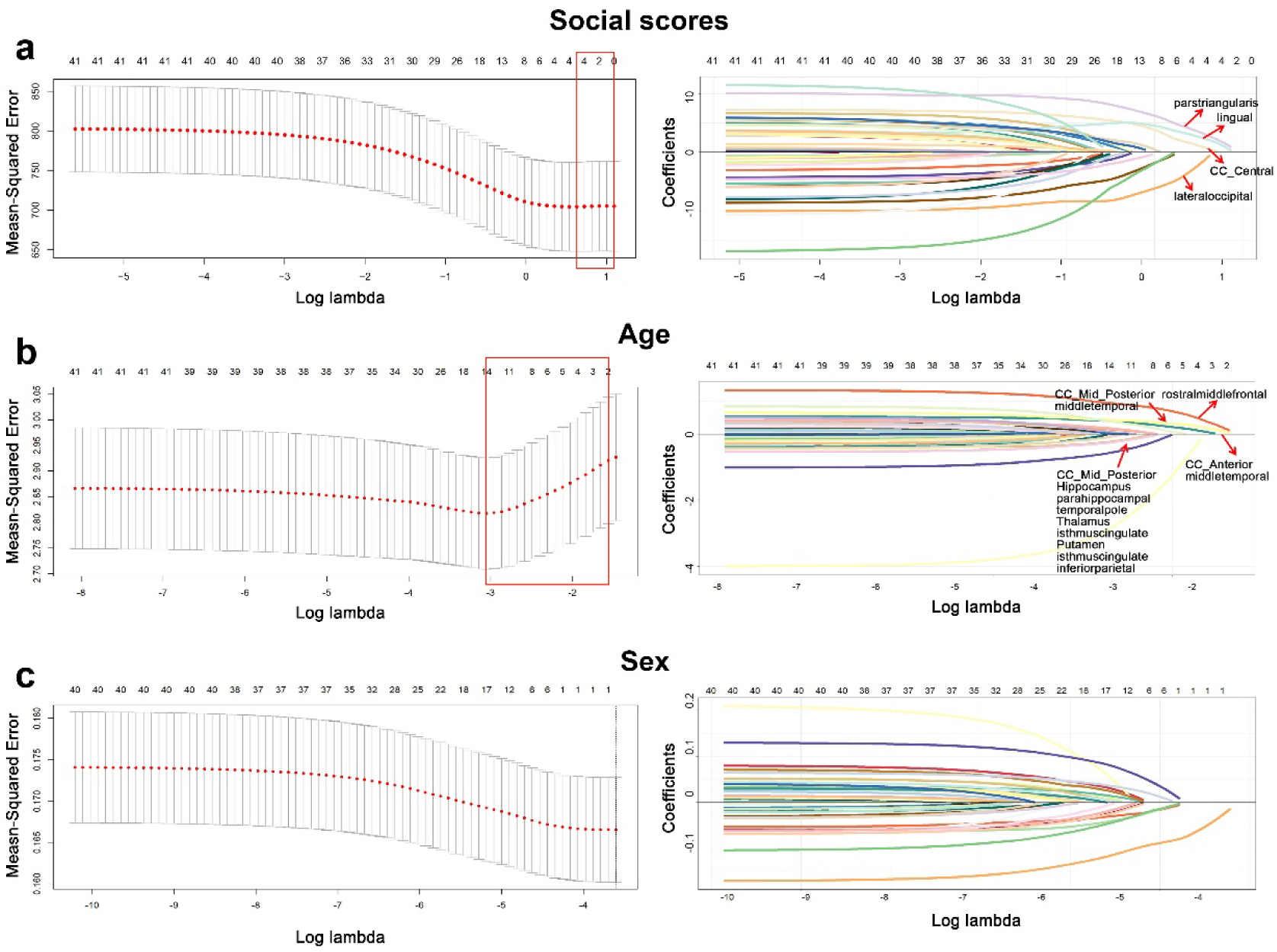
Brain areas that contributed significantly to social cognition deficit scores (a), age(b) and sex(c), based on Glmnet regression analysis. **Left:** Solution paths from the penalized regression with a lasso penalty were used to select the optimal model for the relationship between individualized deviation scores and social cognition deficit scores. The red box highlights the optimal lambda, as determined by cross-validation. **Right:** The complete solution path for all coefficients is shown as a function of the logarithmic tuning parameter, lambda, for the lasso penalty.

**Figure S7.**
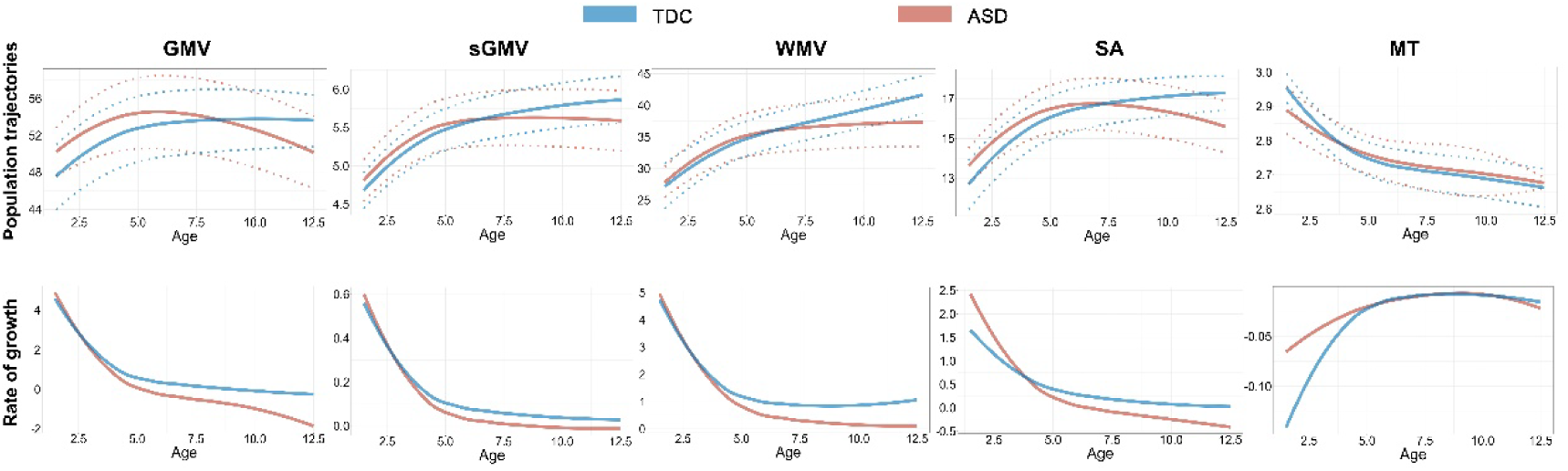
Typical and atypical brain growth chart of females. Top, typical and atypical trajectories of global morphometric phenotype. The median (50% centile) is represented by a solid line, while the 2.5% and 97.5% centiles are indicated by dotted lines. Bottom, rate of growth (the first derivatives of the median trajectory). y axes are scaled in units of the corresponding MRI metrics (10,000 mm^3^ for volume value, 10,000 mm^2^ for surface area and mm for cortical thickness).

**Figure S8.**
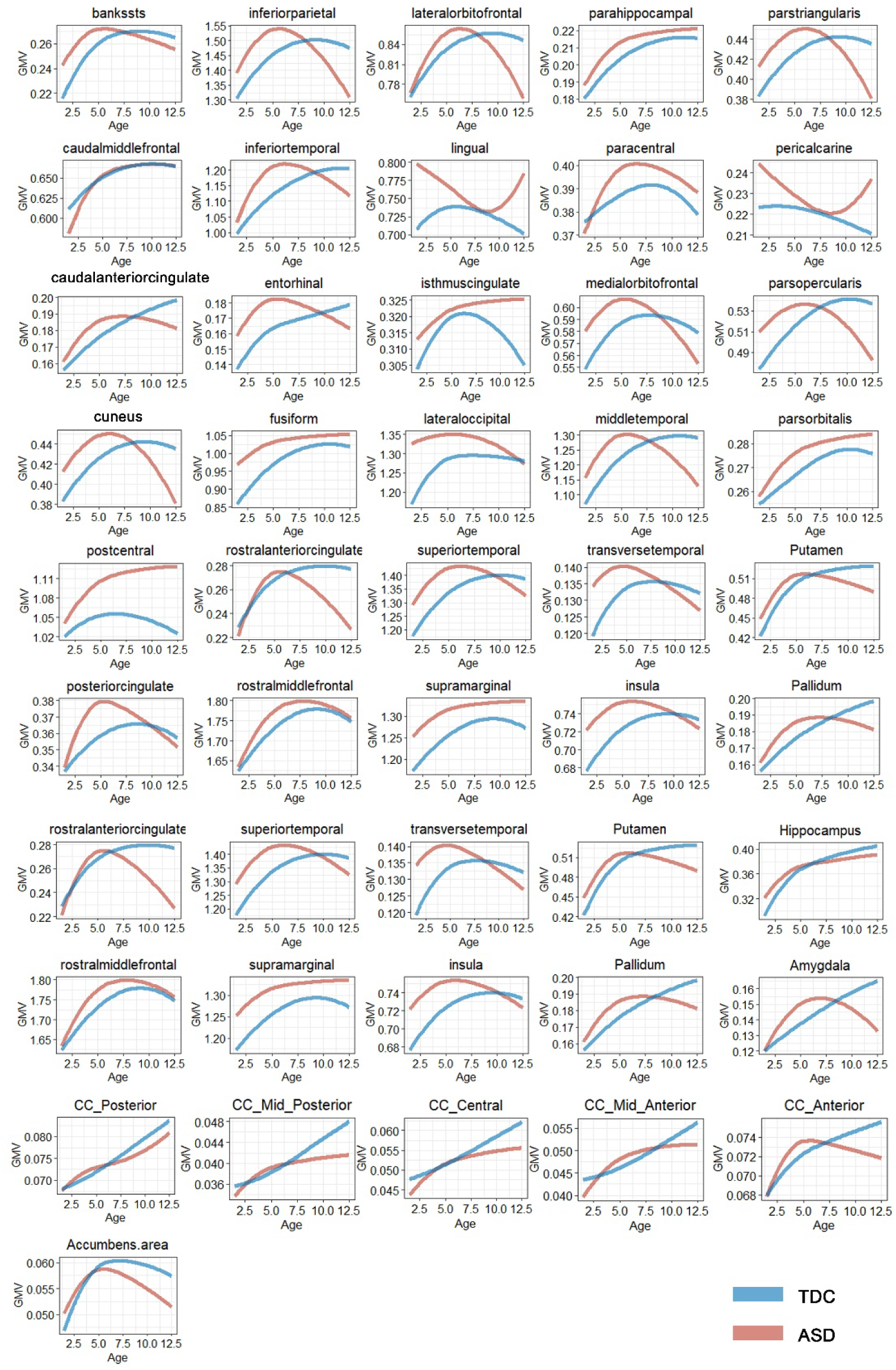
Typical and atypical trajectories of median regional volumes in females for 34 bilateral brain regions as defined in the Desikan-Killiany parcellation and 12 subcortical regions (10,000 mm^3^) These trajectories were fitted to the raw data using the same GAMLSS model used for estimation of global morphometric phenotype trajectories, as shown in Figure S7.

**Figure S9.**
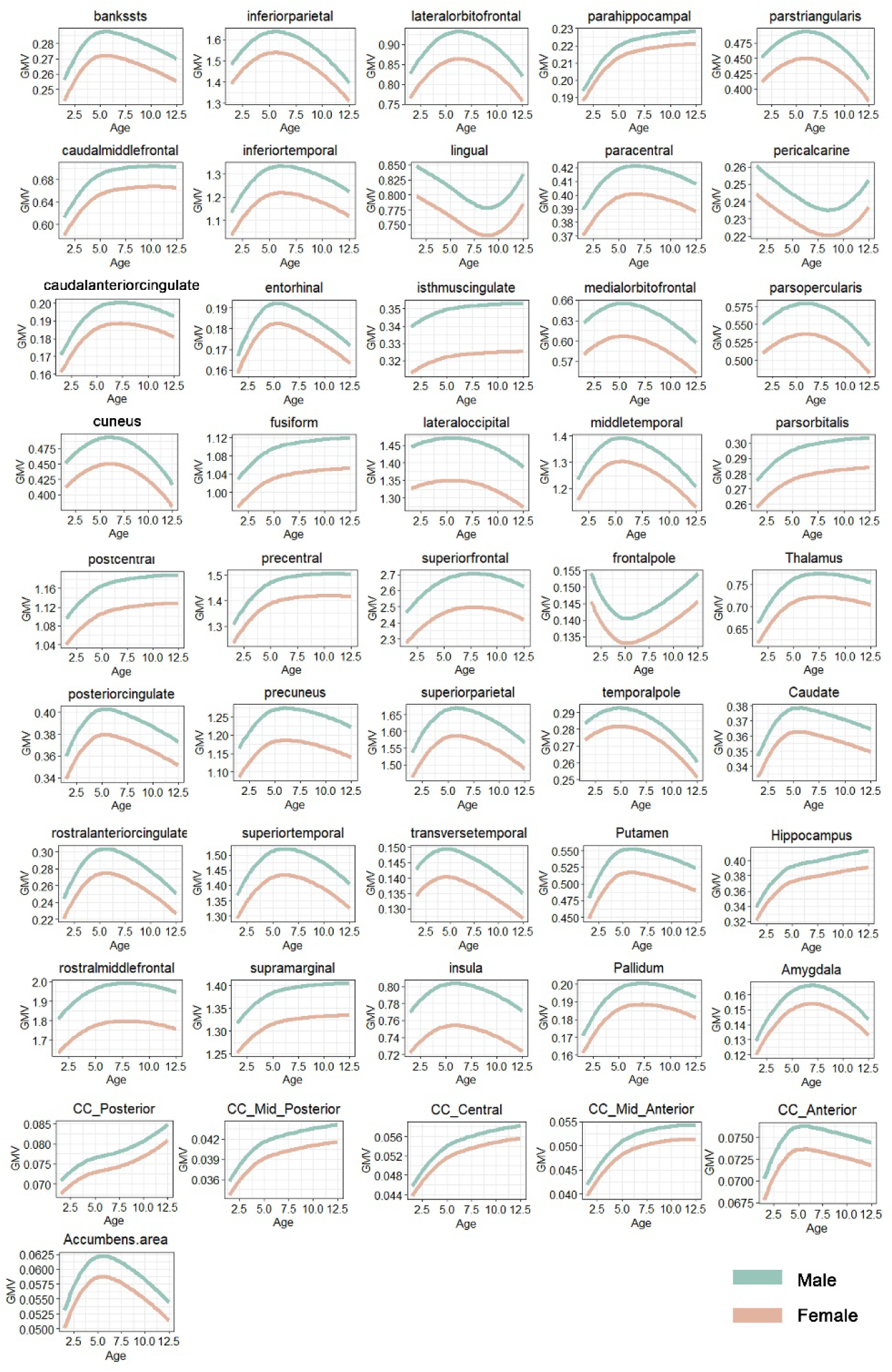
Sex-stratified atypical trajectories of median regional volumes in ASD (10,000 mm^3^)

**Figure S10.**
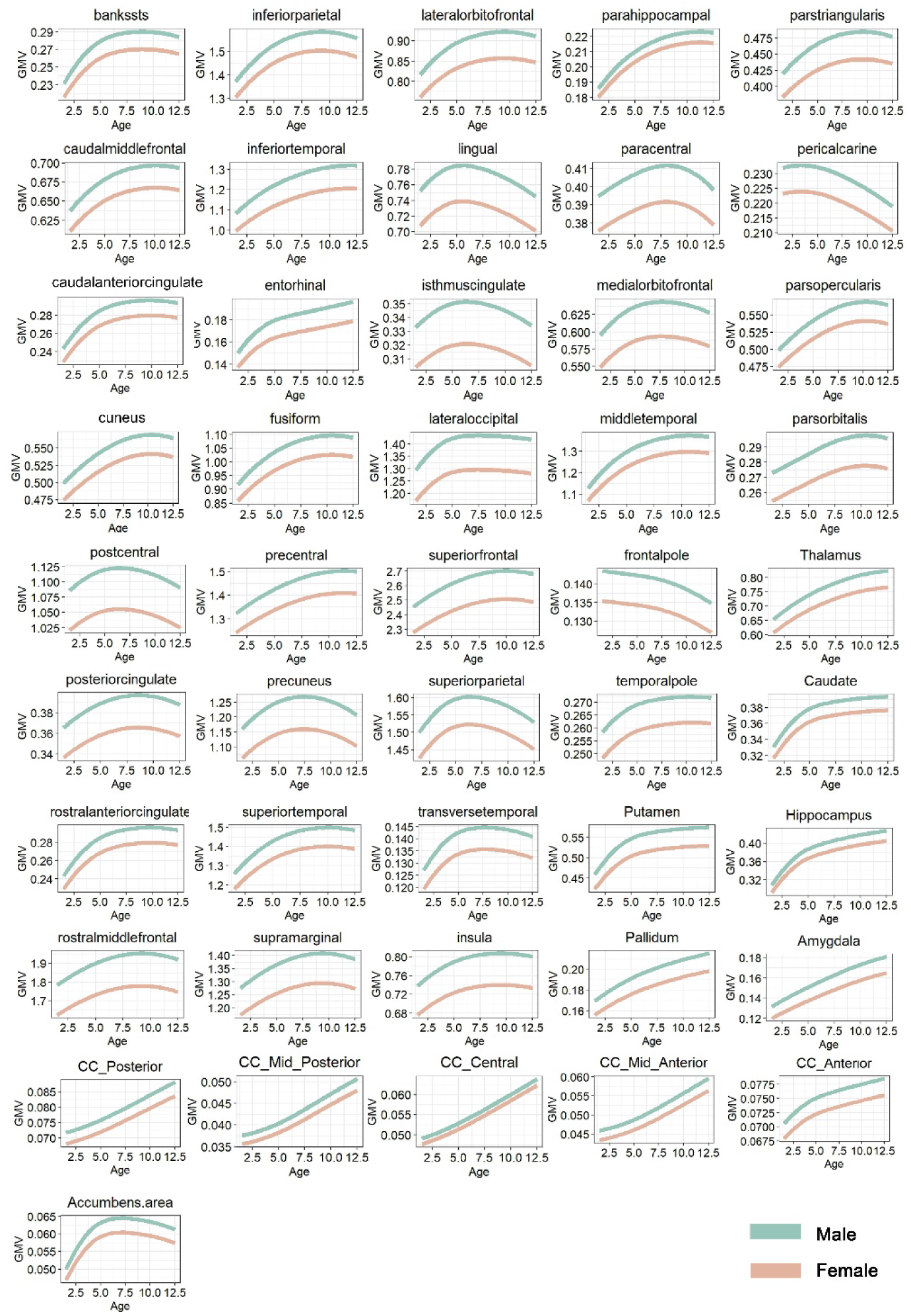
Sex-stratified atypical trajectories of median regional volumes in TDC (10,000 mm^3^)

**Figure S11.**
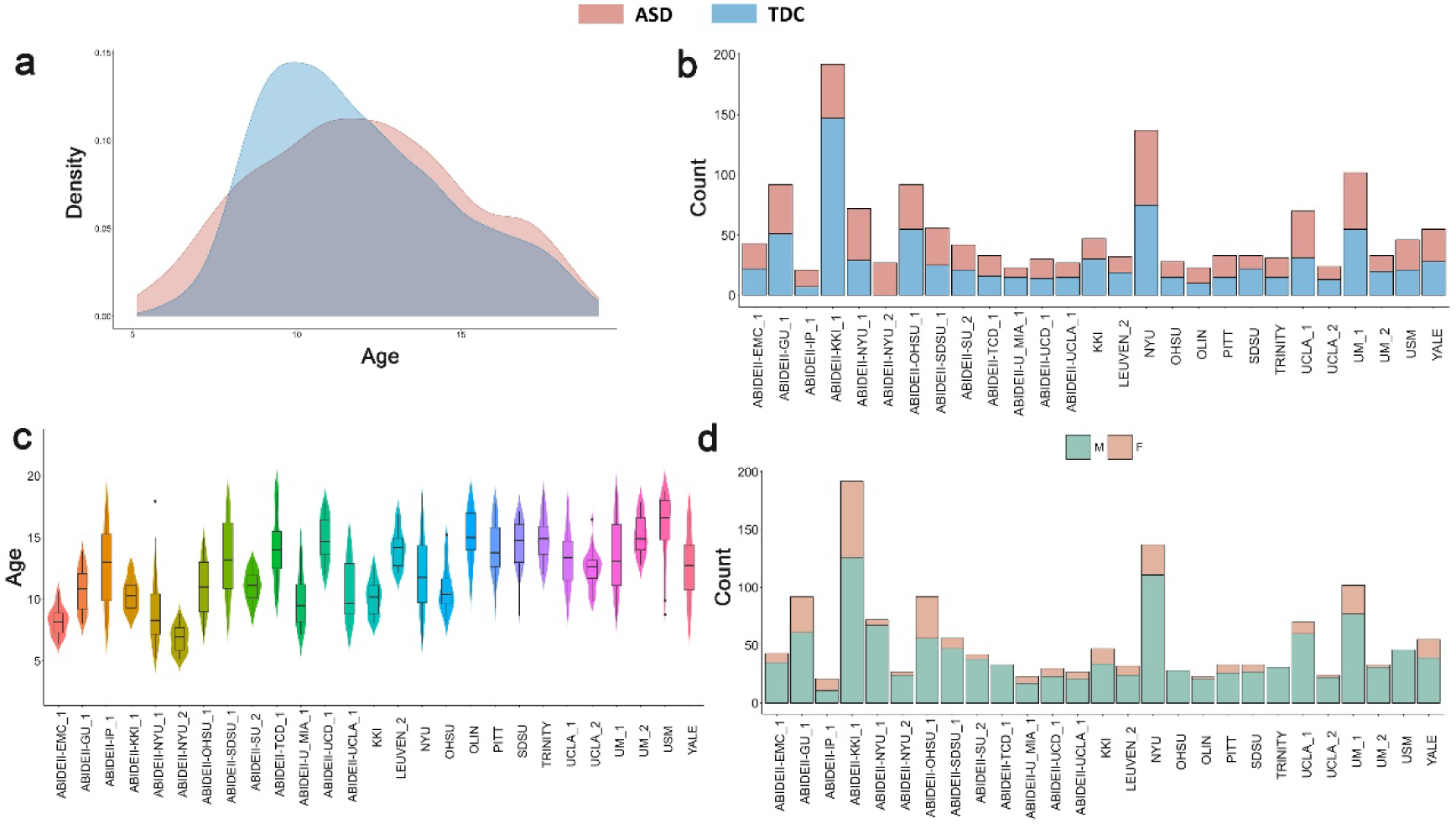
ABIDE Sample Characteristics. **(a)** The distributions of age in ASD and TDC group **(b)** Total number of participants per group for each site. **(c)** Age (in years) for all individuals per site irrespective of diagnostic group. **(d)** Number of males and females for each site irrespective of diagnostic group.

## Supplementary Tables

**Table S1.** Demographics and clinical characteristics of all participants in CABIC.

|  |  | N | AGE<br>(year) | SEX<br>(M/F) | HANDEDNESS<br>(L/R/M) | IQ<br>(N) | DIGNOSIS<br>(N) |
| --- | --- | --- | --- | --- | --- | --- | --- |
| TOTAL | ASD | 1451 | 1.00~12.41 | 1150/301 | 30/837/46 | 286 | 727 |
|  | TDC | 1119 | 1.00~12.92 | 692/427 | 18/763/35 | 259 | / |
| UESTC | ASD | 293 | 1.99~11.00 | 224/69 | 11/272/10 | 37 | 124 |
|  | TDC | 172 | 1.00~12.07 | 115/57 | 3/158/10 | 127 | / |
| JLU | ASD | 312 | 1.00~8.00 | 229/83 | / | 201 | 208 |
|  | TDC | 48 | 1.00~12.9 | 31/17 | / | / | / |
| CHCMU | ASD | 146 | 2.00~6.92 | 407/31 | / | / | / |
|  | TDC | 100 | 1.25~10.25 | 32/68 | / | / | / |
| PLAGH | ASD | 135 | 1.52~12.40 | 108/27 | 0/135/0 | / | / |
|  | TDC | 79 | 3.96~11.00 | 52/27 | 0/135/0 | / | / |
| WXCH | ASD | 111 | 1.82~10.01 | 83/28 | 0/111/0 | 106 | 92 |
|  | TDC | 89 | 1.14~12.80 | 47/42 | 0/45/0 | / | / |
| SYSU | ASD | 109 | 2.15~11.69 | 94/15 | 9/74/26 | 106 | 109 |
|  | TDC | 81 | 5.26~12.64 | 47/34 | 6/68/7 | 81 | / |
| GWCMC | ASD | 79 | 3.05~12.00 | 63/16 | / | / | / |
|  | TDC | 95 | 2.8~9.00 | 59/36 | / | / | / |
| PKU | ASD | 98 | 3.00~5.81 | 87/11 | 3/82/10 | / | 96 |
|  | TDC | 30 | 3.94~5.57 | 24/6 | 0/23/7 | / | / |
| YZU | ASD | 125 | 3.05~12.25 | 111/14 | 0/125/0 | / | 55 |
|  | TDC | / | / | / | / | / | / |
| HMU | ASD | 43 | 2.92~7.94 | 36/7 | 7/36/0 | 37 | 43 |
|  | TDC | 58 | 3.18~8.04 | 48/10 | 3/55/0 | 51 | / |
| BNU | ASD | / | / | / | / | / | / |
|  | TDC | 367 | 4.5~12.9 | 207/160 | 6/355/11 | / | / |

Note: An assessment is considered to include an intelligence quotient (IQ) score if it involves one of the following measures: the Wechsler Intelligence Scale for Children (WISC), Gesell Developmental Schedules (GDS), or the Peabody Picture Vocabulary Test (PPVT). A clinical symptom score is recorded if the assessment includes one of the following: The Autism Diagnostic Interview-Revised (ADI-R), Autism Diagnostic Observation Schedule (ADOS), Childhood Autism Rating Scale (CARS), or Autism Behavior Checklist (ABC).

